# Distractor anticipation during working memory is associated with theta and beta oscillations across spatial scales

**DOI:** 10.1101/2024.10.23.619830

**Authors:** Dennis Y. Jung, Bikash C. Sahoo, Adam C. Snyder

## Abstract

Anticipating distractors during working memory maintenance is critical to reduce their disruptive effects. In this study, we aimed to identify the oscillatory correlates of this process across different spatial scales of neural activity. We simultaneously recorded local field potentials (LFP) from the LPFC and electroencephalograms (EEG) from the scalp of monkeys performing a modified memory-guided saccade (MGS) task. The monkeys were required to remember the location of a target visual stimulus while anticipating distracting visual stimulus, flashed at 50% probability during the delay period. We found significant theta-band activity across spatial scales during anticipation of a distractor, closely linked with underlying working memory dynamics, through decoding and cross-temporal generalization analyses. EEG particularly reflected reactivation of memory around the anticipated time of a distractor, even in the absence of stimuli. During this anticipated time, beta-band activity exhibited transiently enhanced intrahemispheric communication between the LPFC and occipitoparietal brain areas. These oscillatory phenomena were observed only when the monkeys successfully performed the task, implicating their possible functional role in mitigating anticipated distractors. Our results demonstrate that distractor anticipation recruits multiple oscillatory processes across the brain during working memory maintenance, with a key activity observed predominantly in the theta and beta bands.

## 2 Introduction

Imagine the moment just before serving in a tennis match. You plan to serve the ball in a location well beyond your opponent’s reach. With this target in mind, you toss the ball into the air. But just as you enter a service motion, your opponent suddenly moves toward the intended location. This unanticipated movement distracts you, throwing off your timing and, alas, resulting in a faulty serve. Your planned action faltered because your working memory could not withstand the unexpected distraction.

Working memory is a cognitive function that allows for the temporary maintenance and manipulation of information relevant to immediate goal-directed tasks (Baddeley, 1992; D’esposito et al., 1995; Miller and Cohen, 2001; Pasternak and Greenlee, 2005; Miller et al., 2018). It emerges from the coordinated, sustained activation of neurons distributed across the brain capable of transiently encoding task-relevant information (Pasternak and Greenlee, 2005; Christophel et al., 2017; Leavitt et al., 2017; Sreenivasan and D’Esposito, 2019; Dotson et al., 2018; Mejias and Wang, 2022). The Lateral prefrontal cortex (LPFC) plays a central role in maintaining and controlling this information (Goldman-Rakic, 1995; Rainer et al., 1999; Miller et al., 1996, but see Lara and Wallis, 2014; Konecky et al., 2017; Mendoza-Halliday et al., 2024b), as implicated by its enhanced memory-dependent activity (Kubota and Niki, 1971; Fuster and Alexander, 1971; Funahashi et al., 1989) and communication with other relevant brain areas (Salazar et al., 2012; Jacob et al., 2018; Liebe et al., 2012) during the delay period of working memory tasks. In the face of distractors, encoding of memory remains uninterrupted in LPFC (Sakai et al., 2002; Miller et al., 1996; Yoon et al., 2006) and across the brain (Hallenbeck et al., 2021) despite distractors’ detrimental effects on behavior (Brown, 1958; Clapp et al., 2010; Lorenc et al., 2021). While the effects of distractors on working memory are well-documented, less is understood about how anticipating distractors affect memory maintenance and associated neural dynamics across spatial scales of brain signals.

Distractions are thought to be mitigated by either enhancing memory representation or reducing distractor representation, or a combination of both (Foxe and Snyder, 2011; Liesefeld et al., 2020; Lorenc et al., 2021). Effective mitigation often relies on prior knowledge of distractors, such as anticipatory cues or statistcal regularities (Payne et al., 2013; Bonnefond and Jensen, 2012; de Vries et al., 2019; Magosso and Borra, 2024). In the tennis example, if your opponent moves predictably you are less likely to be distracted during your service. Anticipating forthcoming events, like distractions, is known to modulate neural oscillations at different frequency bands and spatial scales, as shown in monkeys, for which prefrontal beta-band oscillations predict anticipation of upcoming stimulus (Wimmer et al., 2016a), and humans, for which scalp alpha-band oscillations reflects suppression of activity when distractors are anticipated (Klimesch et al., 2007; Snyder and Foxe, 2010; Foxe and Snyder, 2011; Bonnefond and Jensen, 2012; Banerjee et al., 2011). Yet, it remains unclear whether these oscillatory patterns associated with anticipatory processes during working memory maintenance are universal across brain signals of different spatial scales. Furthermore, given the role of oscillations in communication between brain areas (Fries, 2005), how these oscillations facilitate communication among brain areas relevant for working memory maintenance in the face of anticipated distractors remains unclear.

In the present study, we examined how distractor anticipation impacts the representation of encoded memory items and the communication between the LPFC and the rest of the brain, we simultaneously recorded local field potential (LFP) from the LPFC and electroencephalogram (EEG) from the scalp of monkeys performing a spatial working memory task, where the monkeys had to remember the location of memory items while ignoring a distracting visual stimulus, which appeared with a 50% probability at a predictable time during the delay period. We found that theta-band oscillations in the LPFC (4 - 8 Hz) and scalp EEG (1 - 6 Hz) were related to working memory dynamics during distractor anticipation through time-series and cross-temporal decoding analyses. Memory-dependent signals in EEG theta-band oscillations initially diminished during the memory delay period, but reemerged after the anticipated time of the distractor when no stimulus was presented. This reactivation of memory was not observed when monkeys failed the task. In the LPFC, theta-band oscillations reflected memory items encoded among LPFC neurons in response to a distractor, which was not observed when the monkeys failed the task. However, unlike EEG theta-band oscillations, no spontaneous reactivation of memory was observed during the anticipated distractor time. We also found that beta-band oscillations (13 - 30 Hz) reflected strong interareal coherence across the brain, with a notable interaction detected between the LPFC recorded intracortically and EEG overlying the occipitoparietal cortex. The strength of beta-band coherence transiently enhanced around the anticipated distractor time, but only in successful trials, implicating its possible functional role in mitigating anticipated distractions.

## 3 Materials and Methods

### 3.1 Subjects

Three adult male macaque monkeys (*Macaca mulatta*), identified as “R” (6 yrs old), “W” (9 yrs), and “T” (8 yrs), were used in this study. All experimental procedures were approved by the University of Rochester Committee on Animal Resources (UCAR) and adhered to the guidelines established by the National Institutes of Health (NIH) *Guide for the Care and Use of Laboratory Animals* (National Research Council, 2011). Surgeries were performed using asceptic technique under isoflurane general anesthesia with perioperative opiate analgesics and antibiotics.

The monkeys were housed either individually or in pairs at the primate facility of the University of Rochester on a 12-hour light-dark cycle (from 7 am to 7 pm) and provided with a nutritionally balanced diet, monitored by on-site veterinarians and animal facility staff. Water intake was regulated, ensuring a minimum of 20 ml/kg/day during experimental sessions and a minimum of 60 ml/kg on non-experimental days. Regular health assessments were performed by trained personnel, including onsite veterinarians and experienced laboratory members familiar with primate care.

Each monkey was surgically fitted with a titanium head post to facilitate head stabilization during experimental sessions. Additionally, two monkeys (R and W) underwent implantation of high-density microelectrode arrays for intracranial neural recordings in the LPFC (area 46v) (Figure 1a). These surgeries were guided based on structure magnetic resonance imaging (MRI) scans obtained with a 3T scanner and intraoperative visual confirmation of the LPFC by identification of arcuate and principal sulci during surgery. Monkey R had a 128-channel NeuroNexus Matrix microelectrode array (NeuroNexus, Ann Arbor, MI, USA) implanted in the left hemisphere. The Matrix array had 16 shanks, each 2 mm long, arranged in a 4 x 4 grid with 400 µm inter-shank spacing; each shank consisted of 8 recording sites spaced 200 µm apart, with the deepest site 38 µm from the tip of the shank. Monkey W received a 96-channel Utah microelectrode array (Blackrock Microsystems, Salt Lake City, UT, USA) in the right hemisphere (Figure 1a). The Utah array consisted of 100 shanks, each 1 mm long, arranged in a 10 x 10 grid with 400 µm inter-shank spacing, with a single recording site at the tip of each shank. Four of the sites were not active (96 active sites). These electrode array implantations were performed once the monkeys learned the task.

**Figure 1:**
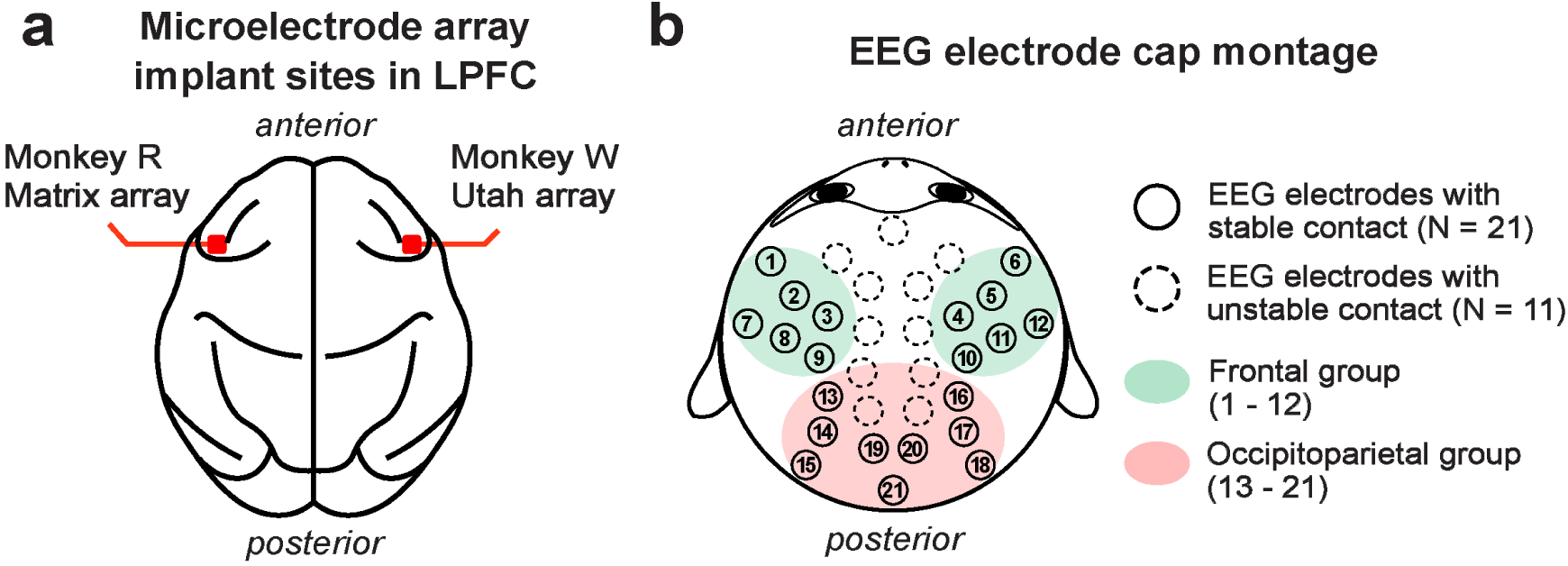
Approximate implant sites for microelectrode arrays used for LFP recording in the LPFC (**a**). Monkey R received a 128-channel NeuroNexus Matrix array in the left LPFC, while monkey W received a 96-channel Utah array in the right LPFC. The 32-channel EEG electrode cap montage (**b**). Electrodes that maintained stable contact with the scalp across all monkeys are marked with filled circles (N = 21 channels). Frontal EEG channels are highlighted in green (N = 12 channels; labeled 1 - 12), and occipitoparietal channels in red (N = 9 channels; labeled 13 - 21).

### 3.2 Behavioral task

Monkeys were trained on a modified version of an memory-guided saccade (MGS) task (Figure 2), a well-established paradigm for studying working memory in primates (Funahashi et al., 1989; Gnadt and Andersen, 1988; Chafee and Goldman-Rakic, 1998; Reinhart et al., 2012). Each trial began with the onset of a circular blue central fixation point (CFP) (RGB: 0, 0, 255; 7 pixels in radius or 0.23 degrees of visual angle (DVA)) at the center of the screen. Once the monkeys successfully fixated on the center fixation point for 500 ms, a target visual stimulus (RGB color: 255,0,255; a 25 pixel radius or 0.81 DVA magenta circle) was briefly presented (50 ms for monkeys R and T, or 100 ms for monkey W; adjusted based on monkeys’ behavioral performance) on one of four possible locations with equal probability (45*^◦^*, 135*^◦^*, 225*^◦^*, and 315*^◦^* relative to the rightward horizontal direction from the CFP; distance from the CFP to the target was 171 pixels or 5.5 DVA).

**Figure 2:**
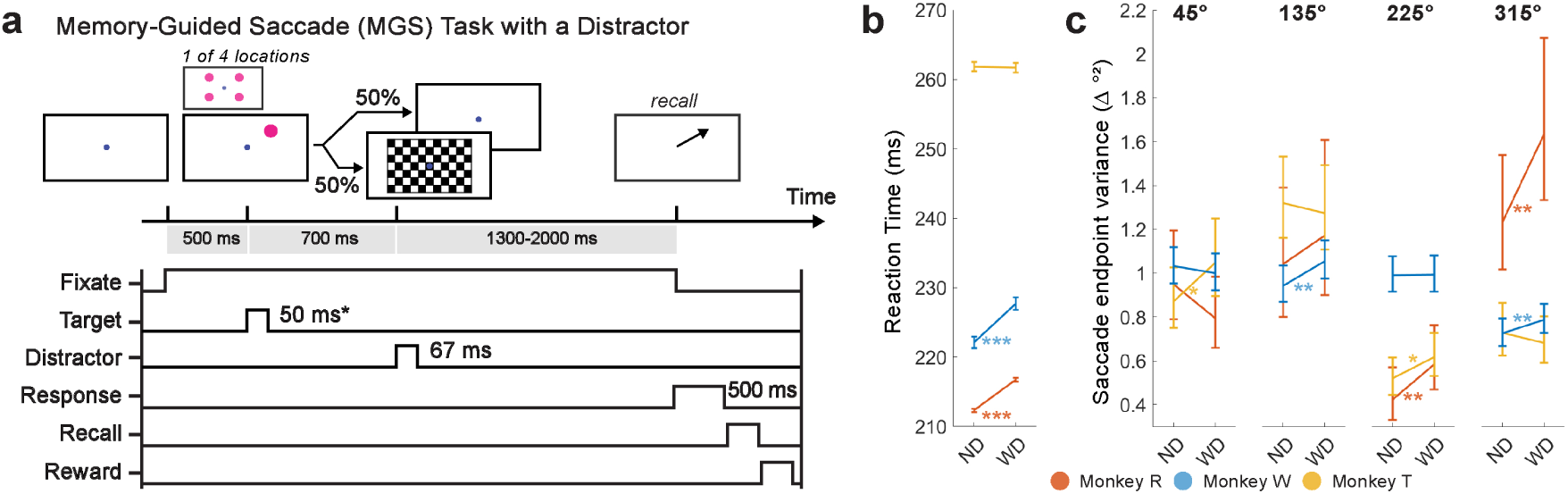
A modified MGS task with a checkerboard stimulus flashed during the delay period (**a**). Each trial began with a 500 ms display of the CFP. After successful fixation on the CFP, a target visual stimulus (item) briefly appeared on the screen (50 ms or 100 ms, depending on monkey). In half of the trials, the checkerboard visual stimulus flashed for 67 ms during the delay period (700 ms relative to the item onset). Monkeys then made a saccade to the remembered location of the item at the end of the delay period to receive a reward. (**b**) Saccadic reaction time (in ms; ±1 SEM). (**c**) Saccade endpoint variance (in Δ *^◦^*^2^; 90% percent bootstrap confidence interval) for each trial and item condition. ND: No Distractor; WD: With Distractor. *: *p* < 0.05, **: *p* < 0.01; ***: *p* < 0.001.

Monkeys were required to remember the location of a briefly-shown target visual stimulus (the “memory item”, or simply “item”) while fixating on the CFP for a ^-^2000-ms delay period. In half of the trials, a checkerboard visual stimulus (a high contrast, contrast-reversing, checkerboard-pattern stimulus; 8 video frames, 8.33 ms per frame, checksize 100 pixels or 3.2 DVA, eight rows and columns) was flashed (67 ms total) during the delay period (700 ms after the target onset). When the CFP disappeared at the end of the delay period, monkeys were required to saccade to within 2*^◦^* of the center of the remembered target location. A correct saccadic response rewarded monkeys with a few drops of water or juice; otherwise, no reward was delivered. The inter-trial interval was fixed at 500 ms when correctly performed; otherwise, the inter-trial interval was longer (1000 ms).

### 3.3 Visual stimuli and experimental setup

Experiments were performed in a sound-attenuated, darkened room. Visual stimuli were generated in MATLAB (MathWorks, Natick, MA, USA) using the Psychophysics Toolbox extensions (Brainard, 1997; Pelli, 1997; Kleiner et al., 2007) and displayed on a gray background (52 *cd/m*^2^) on a gamma-corrected 24-inch ViewPixx monitor (VPixx Technologies, QC, Canada) covered with a transparent acrylic sheet. The viewing distance was approximately 49 cm from the monitor relative to the nasion of the monkey. A digital synchronization output from the ViewPixx monitor was used to synchronize the onset of visual stimuli with neural recordings.

### 3.4 Oculomotor tracking

We used the EyeLink 1000 Plus system (SR Research Ltd., Ottawa, ON, Canada) to track eye position at a sampling rate of 1 kHz. The EyeLink 1000 Plus Camera and EyeLink PM-910 Illuminator Module were positioned above and in front of the monkey’s head, while a coated mirror that transmitted visible light (e.g., from the ViewPixx monitor) and reflected infrared light (e.g., infrared eye tracker from the EyeLink system) was placed at an angle in front of the monkey’s head. For each session, the monkey’s eye position was calibrated using custom experimental control software written in MATLAB and monitored online during the experiment.

### 3.5 Neural recordings

We recorded LFP and EEG at 30 kHz sampling rate using the Ripple Grapevine Neural Interface Processor (NIP) system (Ripple, Salt Lake City, UT, USA).

LFP was recorded from the LPFC of monkeys R and W following a two-week recovery period post-microelectrode implantation surgery to ensure stable LFP acquisition (Figure 1a). It is noteworthy that the Matrix array spans from the superficial to deep layers, while the Utah array is fixed in the superficial layer (1 mm electrode length). As the common cortical reach for these two different arrays was the superficial layer of the LPFC, we focused on analyzing LFP collected from the superficial (1 mm) layers of both arrays. This included LFP collected from 32 channels of the NeuroNexus array in monkey R and 96 channels of the Utah array in monkey W.

For EEG recordings, we utilized a custom 32-channel Sn electrode EEG cap (Electro-Cap International, Inc., Eaton, OH, USA) with a custom montage based on the International 10-20 system (Jasper, 1958). We used a conductive gel (Parker Signagel Electrode Gel; Parker Laboratories, Inc., Fairfield, NJ, USA) to maintain electrical contact with the scalp. Recordings were made from the electrodes that established stable contact with the monkey’s scalp, as some electrodes failed to maintain stable contact due to external head implants, such as a head post. Therefore, out of the 32 available electrodes, 21 electrodes that made stable contact across all monkeys were used for analysis (Figure 1b). Physical EEG electrode positions on the monkey’s head were measured using a Fastrak digitizer (Polhemus Inc., Colchester, VT, USA).

In some sessions, we simultaneously recorded LFP and EEG data from monkeys R (N = 6 sessions) and W (N = 7 sessions).

## 4 Data Analysis

### 4.1 Data preprocessing

LFP and EEG data were processed and analyzed in MATLAB (MathWorks, Natick, MA, USA) primarily using the FieldTrip MATLAB toolbox (Oostenveld et al., 2011) and the Chronux Toolbox (Mitra and Bokil 2008, http://www.chronux.org). Power line noise was removed with a band-stop Infinite Impulse Response (IIR) filter using a MATLAB’s built-in *filtf ilt* function (a 2nd-order two-pass Butterworth filter with a cutoff band of 55 Hz to 65 Hz). We then epoched the data in trials from -500 ms to 2500 ms relative to the onset of the target visual stimulus. The epoched data were high-pass filtered at 0.05 Hz using a 4th-order two-pass Butterworth filter (*ft*_*preprocessing*) then trials and the channels that did not meet our criteria were discarded using the FieldTrip visual artifact rejection function (*ft*_*rejectvisual*). Trials with excessively high z-values (e.g., 3 times the standard deviation), variance, kurtosis, or voltage amplitude were discarded. Trials were then re-referenced (i.e., common averaging re-reference), detrended, and downsampled to 100 Hz. For LFP, we removed the channels with excessively high variance and amplitudes. For EEG, as noted in Materials and Methods, we discarded the channels without stable scalp contact across monkeys (refer to Figure 1b).

### 4.2 Oculomotor movement analysis

To calculate saccadic reaction time, we measured time difference between the time when the CFP disappeared at the end of the delay period of the MGS task and the time when the monkey’s saccade landed within the target location.

To quantify saccade endpoint variance, we computed the generalized variance (GV) of saccadic endpoints for each target location. A saccade endpoint was a two-dimensional gaze position where the monkey’s saccade landed on the target location. Each saccade endpoint was centered by subtracting the centroid of all saccade endpoints corresponding to that target. The GV (unit: *^◦^*^2^) was then quantified by calculating the determinant of the covariance matrix (Wilks, 1932). To test for significant change in saccade endpoint variance between the No Distractor (ND) and With Distractor (WD) conditions, we performed *F* -tests, followed by the calculation of the 90% confidence interval using bootstrapping (iterated 10,000 times).

### 4.3 Power spectral analysis

We identified the dominant oscillatory frequency bands for each subject and neural data set (LFP and EEG). First, we reduced the dimensionality of each neural data set using principal component analysis (PCA) implemented in the FieldTrip *ft*_*componentanalysis* function (parameter: method = “pca”), and focused on the first principal component (PC1), which represents the greatest variance in the data set. To identify significant oscillatory bands, we computed average power spectral density estimates from PC1 using the Chronux *mtspectrumc* function (frequency range: 1 - 12 Hz, 2 Slepian tapers with a time-bandwidth product of 3; frequency range: 12 - 40 Hz: 3 Slepian tapers with a time-bandwidth product of 5) (Slepian and Pollak, 1961). We then characterized the significant oscillatory bands from the average power spectral density estimates using the FOOOF (Fitting Oscillations and One Over F) toolbox, which tests oscillatory (periodic) components against against the aperiodic component (parameters: peak_width_limits = [2, 12], max_n_peaks = 4, min_peak_height = 0.05, peak_threshold = 2.0, aperiodic_mode =“fixed”) (Donoghue et al., 2020).

### 4.4 Time-frequency analysis

To examine LFP and EEG data in both time and frequency domains, we performed time-frequency analysis on individual PC (principal component) trials using a complex Morlet Wavelet Transform (CWT) via our custom MATLAB script. The following parameters were used for CWT: for both LFP and EEG, the number of cycles (*n*) was selected according to the frequency (scale) and was logarithmically increased from 2 to 20 for a frequency range of 2 Hz to 40 Hz at 2 Hz resolution. First, we zero-padded the PC trials by adding a vector of 512 zeros before and after the trial. Then the complex Morlet wavelet (*ϕ*) at each time point, *t*, was calculated for each frequency of interest (*f*):

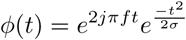

where *j* is the imaginary unit, and the width of the Gaussian (*σ*) depended on the number of cycles for each frequency of interest:

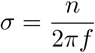

The output of the CWT was first converted to instantaneous power by squaring the output values and then converted to decibel scale relative to the baseline average power computed in the pre-trial period (from -300 ms to -100 ms relative to target onset) and averaged across trials and sessions for visualization.

### 4.5 Time-series decoding

To correlate working memory behavior with oscillatory activity, we decoded the location of the memory item using EEG or LFP oscillatory activity, specifically during the time periods around the target and the distractor onsets.

#### 4.5.1 Filtering data into the oscillatory bands of interest

Before proceeding with decoding analysis, we filtered the processed trials into the frequency bands of interest as identified from our spectral analysis results (Figure 4) using the 4th-order two-pass Butterworth filters via the MATLAB *filtf ilt* function. For EEG, theta-band activity was filtered between 1 Hz and 6 Hz. For LFP, we filtered theta (4 - 8 Hz) and beta bands (13 - 30 Hz), then converted to instantaneous amplitude by taking the absolute values of their Hilbert transformed signals.

#### 4.5.2 Decorrelating LFP and EEG

To reduce the influence of noise sources that were correlated across channels, we applied zero-phase component analysis (ZCA) whitening using the noise covariance matrix estimated from pre-trial activity to decorrelate the feature space in the trial activity of interest (Bell and Sejnowski, 1997; Kessy et al., 2018; Greene and Hansen, 2020; Engemann and Gramfort, 2015). We first calculated the pre-trial whitening matrix *W* from the pre-trial activity that satisfies the following equation:

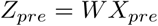

where *W* satisfies the criterion:

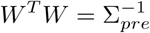

Here, Σ*_pre_* is the noise covariance matrix estimated from pre-trial activity. Then we forced *W* to be symmetrical to satisfy the ZCA criterion:

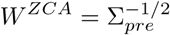

This ZCA whitening matrix, *W ^ZCA^*, was then used to whiten the trial activity of interest, *X*:

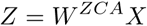

This was performed at the single-trial level. Once all trials were whitened, we normalized *Z* across time and channels by applying a *Z*-score transformation, subtracting the mean and dividing by the standard deviation.

#### 4.5.3 Decoding approach

Before decoding, we first sorted epoched trials into ND and WD conditions. We balanced the number of trials for each trial and item condition (e.g., target item location) by subselecting at random from the larger set to prevent skewed class distribution across conditions. To improve signal-to-noise ratio, we averaged multiple trials within each trial condition and location by sampling 4 trials without replacement, averaging them, and repeating this process until all trials were exhausted (Grootswagers et al., 2017).

We decoded using linear discriminant analysis (LDA) (Hastie et al., 2009). In a preliminary step, we tested and confirmed that LDA produced comparable and faster results when compared to other classification techniques, such as a support vector machine. We performed multi-class categorization using the Error-Correcting Output Codes (ECOC) approach to break down a single multi-class classification task into a series of binary classification tasks (Dietterich and Bakiri, 1994). We used the MATLAB built-in functions: *templateDiscriminant* for LDA and *f itcecoc* for ECOC. We specifically adopted one-versus-all coding (Rifkin and Klautau, 2004), wherein N binary classifiers are trained for N number of classes, which each binary classifier distinguishing one class from the other class. This approach exhibited higher computational performance compared to one-verses-one coding (Nilsson, 1982), as it required less computational time due to the fewer number of binary classifiers to be trained.

Decoding was performed on the averaged trials for each 10-ms time bin (sampling rate: 100 Hz) using the following classification features for each signal type: For EEG decoding, 21 electrode channels with stable scalp contact were used (Figure 1b), and for LFP decoding, microelectrode channels in the superficial 1 mm of the LPFC recording array were used (monkey R: the 32 most superficial channels of the Matrix array; monkey W: all 96 channels of the Utah array). At each time bin, classifiers were trained and tested using an N-fold cross-validation (CV) approach (monkey R, N = 2 folds; monkeys W and T, N = 4 folds). A low N value was chosen for monkey R to mitigate data overfitting during model selection, as fewer total trials were available per recording session.

For a N-fold CV, we partitioned the data set into N non-overlapping trial sets, with each set containing a balanced number of observations selected at random. N-1 sets were randomly chosen to train a classifier and the remainder set was left out to test the classifier. Any excess trials after balancing were omitted. We tested the trained classifier using the MATLAB *predict* function, which returns predicted class labels (i.e., the location of target visual stimulus) based on the held-out data set. The predicted class labels were then compared to the true labels to calculate the classifier’s accuracy, which was determined by dividing the number of correctly classified trials by the total number of trials. Since there were four item locations (classes), the chance-level decoding accuracy was at 0.25 (or 25%), which was subtracted from the actual decoding accuracy. This adjusted decoding performance is referred to as Δ*p*(*decoding*) throughout this paper. We also decoded the items based on their visual hemifield location (left vs. right or upper vs. lower), converting the decoding to binary classification with a chance level of 0.5 (or 50%), which was then subtracted from the actual decoding performance. Then the Gaussian smoothing was applied using the MATLAB *smoothdata* function (parameters: method = “gaussian”, window = 3). Once we evaluated the classifier performance, we repeated this N-1 more times to complete the CV. The entire decoding procedure, starting from the data preparation step of averaging every 4 randomly selected trials, was repeated 20 times to improve robustness when rigorously testing different combination of averaged trials during CV. Additionally, we decoded incorrectly performed trials by evaluating them using classifiers trained on correct trials.

#### 4.5.4 Isolating the contributions of frontal and occipitoparietal EEG activity during decoding

To test how different brain areas contribute to working memory maintenance during anticipated distraction, we re-trained and cross-validated classifiers using subsets of EEG electrodes grouped by their position: (1) the frontal EEG channel group, which included 12 electrodes channels (EEG channels 1 - 12, Figure 1b), and (2) the occipitoparietal EEG group, which included 9 electrode channels (EEG channels 13 - 21). Using each group of EEG channels as classifier features, we performed the same LDA-based decoding analysis described earlier.

#### 4.5.5 Cross-temporal generalization

To test similarity in oscillatory activity patterns across time bins, we employed cross-temporal generalization (King and Dehaene, 2014; Stokes, 2015; Grootswagers et al., 2017) (Figure 3). This method was built upon the decoding procedure described earlier, but instead of evaluating the classifiers only at the trained time bin, each classifier was also tested across all other time bins. If the classifier significantly above chance at different time bins, it indicates that the classifier is generalizable and that the underlying oscillatory activity patterns encoding the memory item are similar across these time bins.

**Figure 3:**
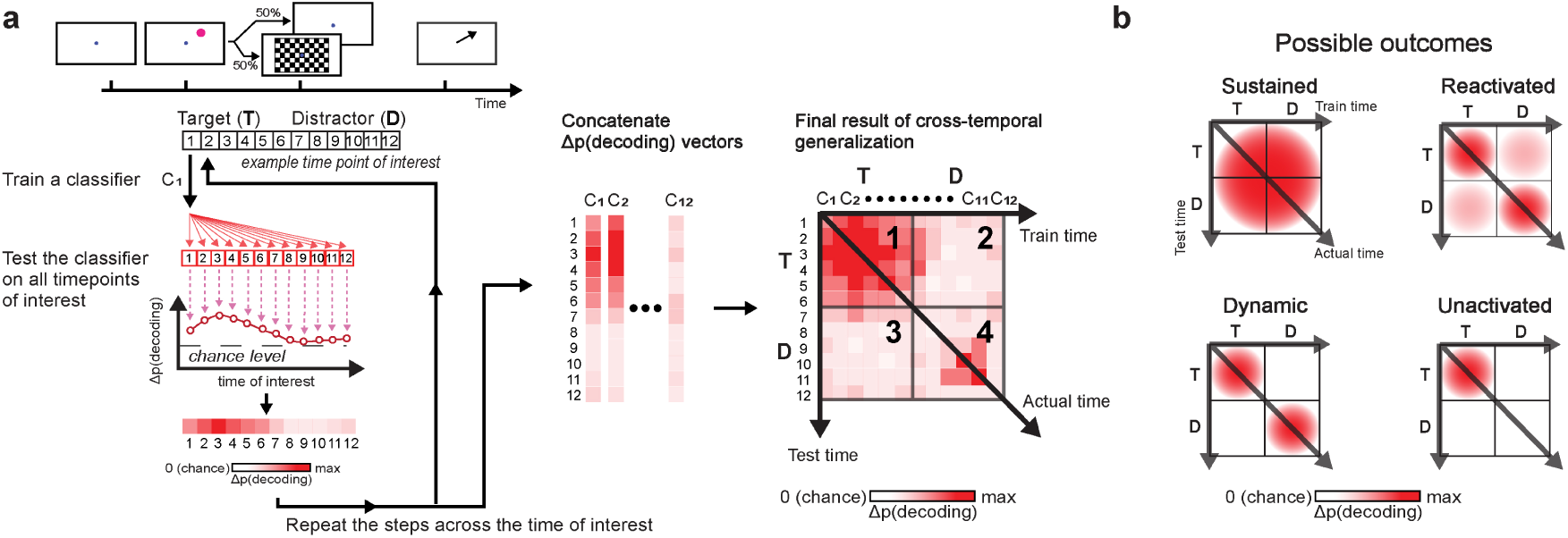
Cross-temporal generalization schematic (**a**). Decoding was performed at each time bin by training a classifier (e.g., C_1_) at one time bin (e.g., t = 1) and evaluating it across all time bins of interest (e.g., 1 ≤ t ≤ 12). This generated a vector of decoding accuracies for each classifier. Repeating this process across all time bins produced vectors of decoding accuracies that were concatenated into a matrix to complete the analysis. In the final result of cross-temporal generalization, panel 1 shows generalization within the post-target period; panels 2 and 3 show generalization between the post-target period and the post-anticipated-distractor/post-distractor period; panel 4 shows generalization within the post-anticipated-distractor/post-distractor period. Color intensity reflects decoding accuracy, Δp(decoding), with values adjusted by one over total number of classes (e.g., subtracted 1/4, when decoding based on the 4 item locations). Possible outcomes (**b**) include: sustained, reactivated, dynamic, and unactivated. “Sustained” indicates consistent generalization over time, reflecting no significant change in the underlying neural activity. “Reactivated” refers to common neural activity patterns at two distinct time periods, though not continuously. “Dynamic” indicates that the underlying neural activity patterns differ between distinct time periods. “Unactivated” occurs when no encoding of items is observed at stimulus onset, here specifically at distractor onset.

#### 4.5.6 Statistical analyses of decoding accuracy

We compared the mean decoding accuracy (Δ*p*(*decoding*)) against the mean of trial-shuffled decoding accuracy (chance-level performance, around 0) using cluster-based two-sample, one-tailed permutation tests (Maris and Oostenveld, 2007; Groppe et al., 2011), as follows. First, the trial-shuffled decoding accuracy was calculated by running the same decoding analysis explained previously but with class labels shuffled across trials. After *t*-testing each time bin (*α* = 0.05), we identified cluster of contiguous bins with significant *t* scores and summed these scores to obtain cluster-level *t* masses. We then compared these *t* masses between the actual and shuffled decoding performances to examine significance (e.g., cluster *α* = 0.05). This same procedure was followed for all cluster-based permutation tests described in this report (i.e., for testing coherence and Granger causality). For comparing between trial conditions (e.g., ND vs. WD), we employed cluster-based two-sample, two-tailed permutation tests.

#### 4.5.7 Visualizing activation patterns using the classifier weights

To visually depict the contributions of individual EEG electrode channels on decoding performance, we constructed topographical maps based on the classifier weights of the trained classifiers. Classifier weights themselves, however, need not necessarily implicate the actual importance of the neural activity observed at the corresponding channels, because the variance of individual channels differs. For example, electrode channels detecting large voltage amplitude changes can result in small classifier weights whereas those with small voltage amplitude can result in large classifier weights. To account for this, we instead calculated the activation pattern by multiplying the classifier weights with the covariance matrix of the training data (Haufe et al., 2014; Fahrenfort et al., 2017). The activation pattern was then normalized across electrodes by applying a *Z*-score transformation, subtracting its mean across electrodes and dividing by its standard deviation across electrodes. Then *Z*-scored activation patterns were averaged across all EEG recording sessions for depiction.

### 4.6 Coherence analysis

To test the strength of functional connectivity with respect to the LPFC, we estimated coherence between the dimensionally-reduced LFP from the LPFC and EEG at each included EEG electrode channel. To reduce noise and dimensionality, we first applied PCA to the LFP data, then analyzed the PC1. For the EEG data, we improved topographical localization and reduced volume conduction by applying the surface Laplacian transform using the FieldTrip *ft*_*scalpcurrentdensity* function (parameters: method = ‘finite’, degree = 7) (Oostendorp and van Oosterom, 1996; Huiskamp, 1991). After the pre-processing, we used the Chronux *coherencyc* function (parameters: tapers = [5, 7], pad = 1) to calculate coherence between the PC1 of the LPFC LFP and the EEG signals at each EEG electrode channel.

Brain areas may appear functionally connected if they independently respond to task stimuli or contexts within the same oscillatory band. Therefore, we utilized a more stringent measure called “excess coherence” (Δcoherence), which adjusts the original coherence estimate (Snyder et al., 2018). Excess coherence was calculated by subtracting coherence by the average of 200 trial-shuffled versions, where the trial order of the EEG data set was randomly permuted for each iteration. We tested for excess coherence significantly greater than zero using a cluster-based one-sample, one-tailed permutation test. Throughout this paper, all reported coherence values are excess coherence.

Coherence was computed using different time windows: longer windows were used to visualize topographical maps of coherence, while shorter windows were taken to examine the temporal dynamics of coherence. To visualize overall functional connectivity on topographical maps, we calculated coherence during the first second of the task (0 ms - 1000 ms relative to target onset) for each recording session and then averaged across sessions. This analysis was done separately for each monkey, as the microelectrode arrays were implanted in different LPFC sites –one in the left hemisphere and another in the right hemisphere –but both still in the area 46v. We also examined the temporal dynamics of coherence across trial conditions, specifically testing whether functional connectivity differed when monkeys remembered memory items contralateral (CONTRA) versus ipsilateral (IPSI) to the LPFC implant site. Before estimating coherence, trials were grouped according to these conditions. Coherence was then computed in a 500-ms window with a 250-ms overlapping sliding step. Statistical significance of coherence was examined using a cluster-based one-sample, one-tailed permutation test against 0, as the coherence values had already been subtracted from their mean trial-shuffled counterparts.

#### 4.6.1 Granger causality analysis

To examine the directed functional interactions between the LFP and EEG, we performed Granger causality analysis (Granger, 1969; Geweke, 1982). We first computed Fourier spectra of LFP PC1 and EEG using the Chronux *mtspectrumc* function (parameters: fpass = [1,40], tapers = [5,7], trialave = 0, pad = 1). These spectra were then input into the FieldTrip *ft*_*connectivityanalysis* function (parameters: method = “granger”, granger.conditional = “no”, granger.sfmethod = “bivariate”) to calculate Granger causality index (GCI). For significance testing of GCI between directions, we used a cluster-based two-sample, two-tailed permutation test comparing the computed GCI to its trial-shuffled counterparts (i.e., chance-level GCI), using the same procedure described in coherence analysis.

## 5 Results

We investigated how oscillatory activities relate to distractor anticipation during working memory maintenance. We tasked monkeys with a modified MGS task, where a distracting visual stimulus (i.e., checkerboard) was presented in half of the trials at a fixed time during the delay period. To assess behavior, we measured the effect of the distractor on saccadic reaction time and saccade endpoint variance. For neural signals, we analyzed LFP recorded from the LPFC and EEG from the scalp, testing how oscillations in these signals were associated with distractor anticipation and working memory maintenance.

### 5.1 Distractor negatively affected behavioral performance

Overall, monkeys performed the task well, as shown by good rates of correctly reporting the location of the remembered item. Total hit rates (correct trials out of correct trials and incorrect trials) for each monkey were 72.8% (2829 of 3886 trials), 95.7% (16347 of 17084 trials), and 80.8% (4944 of 6118 trials) for monkeys R, W, and T, respectively. Mean hit rates across sessions for monkeys R, W, and T were 72.1 ± 3.8% (mean ±1 SEM; N = 8 sessions), 96.5 ± 0.4% (N = 14 sessions) and 80.2 ± 2.3% (N = 8 sessions), respectively. In addition, there was no significant difference in hit rates with or without the presentation of the checkerboard stimulus (two-sample *t*-test, two-tailed; monkeys R, W, and T: *p* = 0.77, 0.47, and 0.17, respectively) (Supplementary Figure 1). These behavioral results suggest that the monkeys were well-trained and showed consistent performance across recording sessions.

Despite generally good performance on the task in the presence of the checkerboard stimulus, we found two signatures of its distracting effect on working memory performance (Figure 2b,c): (1) slowing of saccadic reaction time and (2) increased saccade endpoint variance.

Saccadic reaction times slowed in monkeys R and W when the checkerboard stimulus was presented (WD, monkey R: 227.7 ± 0.8 ms, N = 737 trials; monkey W: 216.7 ± 0.3 ms, N = 8273 trials), as compared to when it was absent (ND, monkey R: 222.1 ± 0.9 ms, N = 718 trials; monkey W: 212.3 ± 0.2 ms, N = 8122 trials; Wilcoxon rank-sum test, two-tailed; monkey R: *p* < 0.001, *z* = -4.71 ; monkey W: *p* < 0.001, *z* = -14.1). Reaction times were, however, not different between conditions for monkey T (ND, 261.8 ± 0.7 ms, N = 2178 trials; WD, 261.7 ± 0.7 ms, N = 2165 trials; Wilcoxon rank-sum test, two-tailed; *p* = 0.98, *z* = -0.0279), suggesting individual variability in response.

The checkerboard stimulus generally increased saccadic endpoint variance, reflecting increased error during memory recall (Figure 2c). In monkey R, it significantly increased the variance for target visual stimuli presented at 225° (*F*_(260,231)_ = 1.38, *p* = 0.006) and 315° (*F*_(337,328)_ = 1.32, *p* = 0.005). However, there was no significant increase for stimuli at 45° (*F*_(350,354)_ = 0.84, *p* = 0.95) or 135° (*F*_(207,240)_ = 1.13, *p* = 0.18). In monkey W, the variance was significantly increased at 135° (*F*_(2202,2188)_ = 1.12, *p* = 0.005) and 315° (*F*_(2168,2197)_ = 1.09, *p* = 0.025), while there was no significant increase at 45° (*F*_(2218,2205)_ = 0.97, *p* = 0.78) or 225° (*F*_(2183,2180)_ = 1.00, *p* = 0.47). In monkey T, the variance was significantly increased at 45° (*F*_(586,600)_ = 1.20, *p* = 0.012) and 225° (*F*_(607,605)_ = 1.19, *p* = 0.016), but not at 135° (*F*_(609,615)_ = 0.96, *p* = 0.68) or 315° (*F*_(598,604)_ = 0.94, *p* = 0.79). Thus, for all three monkeys, the presence of the checkerboard led to increased saccade endpoint variance for two of the target locations, while the other two locations showed no difference.

These behavioral metrics demonstrated that the checkerboard stimulus during the task acts a salient distractor that captures attention and hinders overall working memory performance.

### 5.2 Oscillatory characteristics in monkey EEG and LPFC LFP

#### 5.2.1 Theta and beta power predominates during working memory maintenance

Because we were interested in the role of neural oscillations in working memory, we first sought to characterize the center frequencies, bandwidths and dynamics of the oscillatory activity of our recordings during our spatial working memory task.

We identified dominant oscillations during the MGS task by examining the spectral characteristics of LPFC LFP and scalp EEG signals. Overall, theta- and beta-band power was prominent in both signals as shown from power spectral analysis (Figure 4). In LFP, the mean center frequency of the theta band across monkeys was 4.8 ± 1.8 Hz (±1 SEM), with a mean bandwidth of 2.1 ± 2.7 Hz (±1 SEM) (Figure 4c,d; 1-12 Hz). The beta band had a mean center frequency of 25.0 ± 4.5 Hz and a mean bandwidth of 6.6 ± 2.7 Hz (Figure 4c,d; 12-40 Hz). In EEG, the theta band had a slightly lower mean center frequency of 4.4 ± 0.7 Hz and a mean bandwidth of 1.8 ± 0.5 Hz (Figure 4j,k; 1-12 Hz), while the beta band had a mean center frequency of 24.3 ± 5.0 Hz and a mean bandwidth of 8.3 ± 3.8 Hz (Figure 4j,k; 12-40 Hz), closely aligning with the LFP beta band range. However, no dominant alpha-band power (8 - 13 Hz) was observed in either signal type.

**Figure 4:**
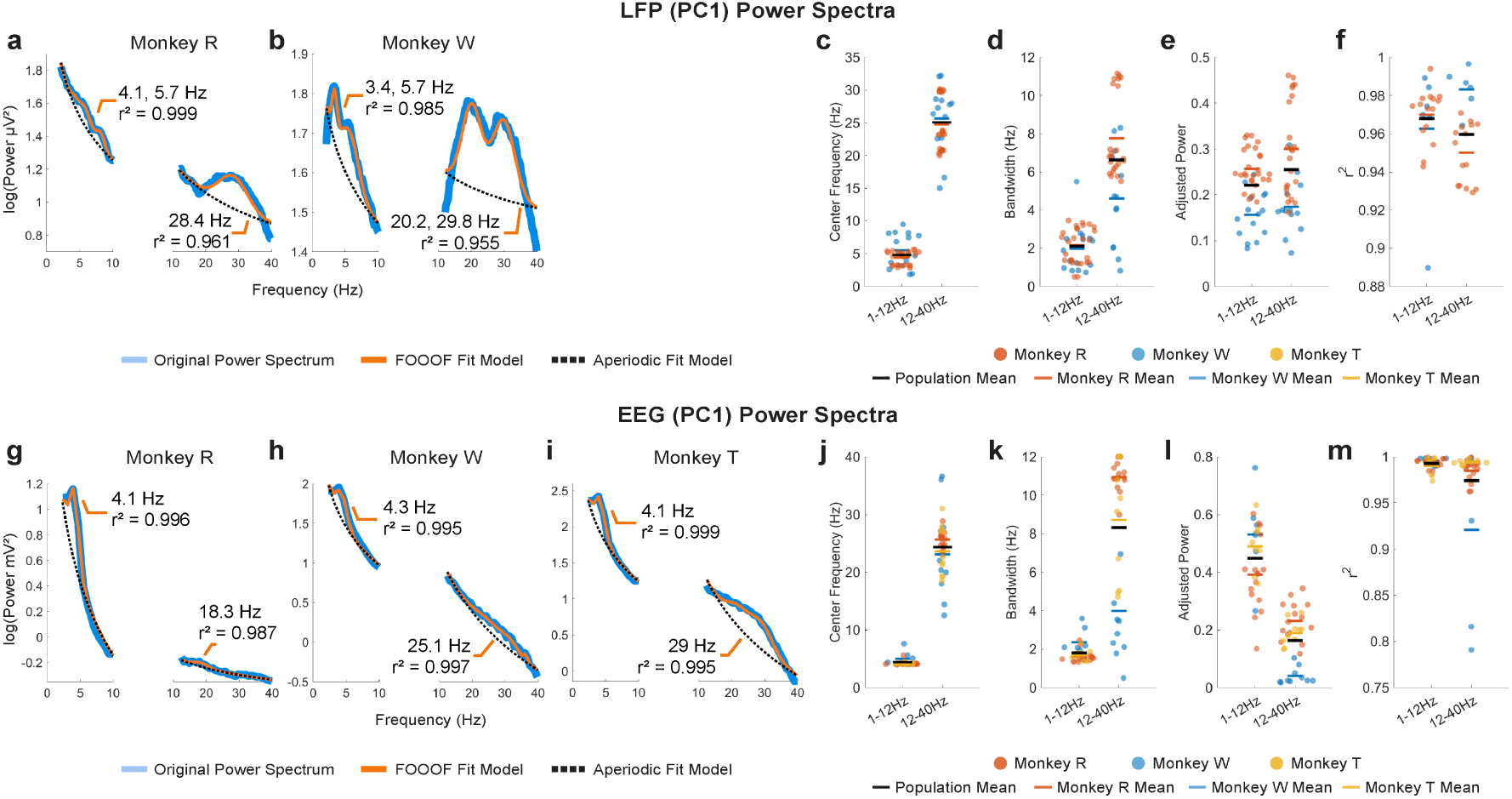
Representative examples of PC1 power spectra (blue traces) and their FOOOF model fits (orange traces) for LPFC LFP in monkeys R (**a**) and W (**b**) and for EEG in monkeys R (**g**), W (**h**), and T (**i**), with aperiodic fits (dotted black traces). FOOOF results for LFP –center frequency (**c**), bandwidth (**d**), adjusted power (**e**), and goodness-of-fit (*r*^2^) (**f**) –are shown for monkeys R and W. For EEG, the FOOOF results –center frequency (**j**), bandwidth (**k**), adjusted power (**l**), and goodness-of-fit (**m**) –are shown for monkeys R, W, and T. Population mean of all monkeys shown in horizontal black bars.

We also examined the dominant spatiotemporal dynamics of the neural signals by applying time-frequency analysis to dimensionally reduced the signals using PCA (Supplementary Figure 2). In the first principal component (PC1) of the LFP, both monkeys exhibited robust power (estimated in decibels relative to baseline) stretching from theta (4 - 8 Hz) to beta (13 - 30 Hz) bands in response to target or distractor stimulus but none in the absence of stimuli (Supplementary Figure 2a,c). In the second principal component (PC2) of the LFP, only monkey W showed beta-band activity that persisted beyond the anticipated distractor onset (700 ms; Supplementary Figure 2d, ND). This activity was briefly disrupted by the distractor presentation but recovered before fading around 1200 ms. Significant differences between trial conditions (ND vs. WD were only found after the distractor was presented (Supplementary Figure 2a-d). On the other hand, the time-frequency representations of the EEG PC1 showed subtle but distinct low theta-band activity (1 - 6Hz) consistently across all monkeys, regardless of distractor presence, while no noticeable beta-band changes were detected (Supplementary Figure 2e,g,i).

Taken together, we concluded that the dominant oscillatory activity during the working memory task occurs in the theta band (LFP: 4 - 8 Hz, EEG: 1 - 6 Hz) and the beta band (LFP: 13 - 30 Hz).

### 5.3 EEG theta-band oscillations in working memory maintenance and distractor anticipation

Across spatial scales, theta-band oscillations were predominant during the task, aligning with prior research on their possible roles in working memory (Raghavachari et al., 2001; Itthipuripat et al., 2013; Riddle et al., 2020). Given that the activity is predominant during working memory task and appears to modulate during task events as depicted from our time-frequency analysis, we hypothesized that scalp EEG theta-band activity reflects the information associated with the memory item during the task. We also tested how presence of an anticipated distractor modulated the theta-band activity. To test these hypotheses, we conducted decoding and cross-temporal generalization analyses on continuous EEG theta-band activity, focusing on the period when the distractor appeared with 50% probability during the delay. By comparing the results between correct and incorrect trials, we attempted to deduce the information reflected by this activity.

#### 5.3.1 Theta-band activity at anticipated distractor time predicted task performance

We first tested whether EEG theta-band activity was associated with the memory item by decoding its location in 10-ms time bins (because data were downsampled to a 100 Hz sampling rate) throughout the task period.

When the monkeys performed the task correctly, we reliably decoded the location of the memory item after its brief presentation on the screen, with peak decoding accuracy occurring about 100 ms after target onset (peak Δ*p*(*decoding*) = 0.13, i.e., 13 percentage points above chance; 1 cluster, *p* < 0.001, cluster-based permutation test with *α <* 0.05*, cluster α <* 0.05; N = 29 sessions combined across monkeys; Figure 5b). However, this significant decoding did not persist through the delay period, gradually receding to chance-level performance. We also decoded the memory item following either the anticipated distractor onset (i.e., ND condition) or the actual distractor onset (i.e., WD condition). In both conditions, significant decoding occurred a few hundred milliseconds after these onsets (Figure 5c,d), as though the encoding strength of memory items increases after the anticipated distractor time. Significant decoding appeared earlier and lasted longer when distractor did not appear (ND: peak Δ*p*(*decoding*) = 0.04 at 1000 ms, lasting 240 ms; 1 cluster, *p* < 0.001) compared to when it actually appeared (WD: peak Δ*p*(*decoding*) = 0.03 at 1020 ms, lasting 50 ms; 1 cluster, *p* < 0.001). Although the peak decoding occurred around the same time (ND: 1000 ms; WD: 1020 ms) for both conditions, significant decoding was delayed about 100 ms when the distractor was shown (ND: 900 ms; WD: 1010 ms), showing that the distractor briefly disrupted memory encoding.

**Figure 5:**
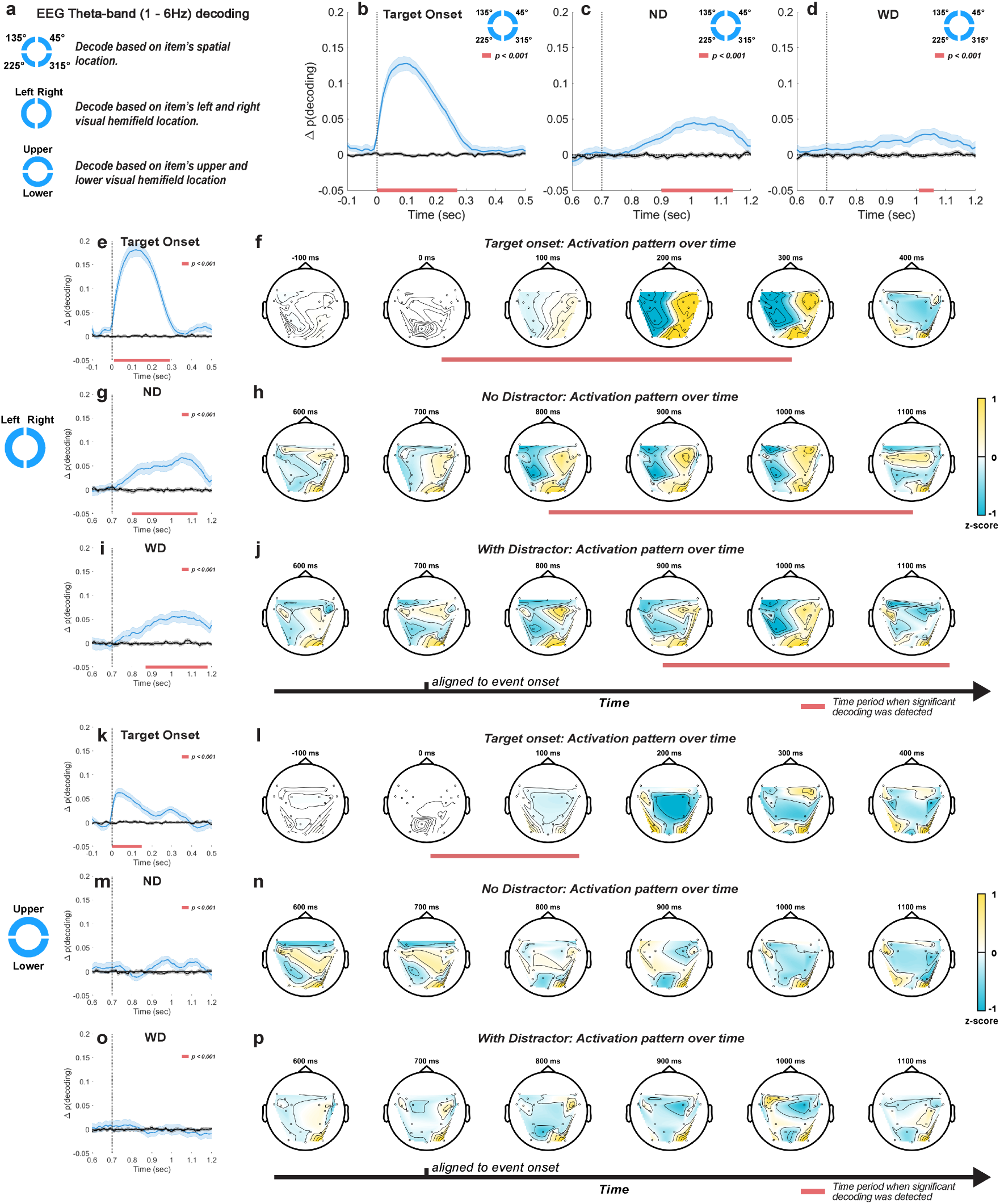
Decoding approaches taken for EEG theta-band (1 - 6 Hz) activity (**a**). Location-based decoding for the post-target (**b**), post-anticipated-distractor (ND) (**c**), and post-distractor (WD) (**d**) periods. Left-versus-right visual hemifield decoding for the post-target (**e**), post-anticipated-distractor (ND) **g**, and post-distractor (WD) (**h**) periods, with their corresponding time-resolved activation patterns (**f**, **h** and **j**, respectively). Upper-versus-lower visual hemifield decoding for the post-target (**k**), post-anticipated-distractor (ND) (**m**), and post-distractor (WD) (**o**) periods, with their corresponding time-resolved activation patterns (**l**, **n** and **p**, respectively). Mean decoding accuracy (blue traces, ±1 SEM) and their trial-shuffled results (black traces, ±1 SEM), which represent chance-level performance. Horizontal red bars at the bottom of each plot indicate significant decoding clusters (*p* < 0.001, N = 26 sessions, combined across monkeys). Activation patterns normalized by *Z*-score transformation.

To test whether the spontaneously increased decoding accuracy after the anticipated distractor time reflects memory-related signals, we examined incorrectly performed trials, where the monkeys failed to make a saccade to the correct location by evaluating classifiers trained on correct trials with incorrect trials. Compared to correctly performed trials, we found lower decoding performance at target onset, reflecting a reduced representation of memory items (peak Δ*p*(*decoding*) = 0.06 at 100 ms, lasting 230 ms, 1 cluster, *p* < 0.001; Supplementary Figure 4a). Furthermore, no significant decoding was seen during either the anticipated or actual distractor periods (ND and WD; Supplementary Figure 4a). These results suggest that EEG theta-band activity reflected the information about memory items following the anticipated distractor time only when the monkeys correctly performed the task.

#### 5.3.2 Theta-band oscillations predominantly reflected left versus right visual hemi-field location

Previous studies implied that the encoding of visual items relies heavily on brain areas contralateral to the location of the item displayed in the visual field (Funahashi et al., 1989; Sawaguchi and Iba, 2001; Pasternak and Greenlee, 2005; Hagler Jr and Sereno, 2006; Wimmer et al., 2016b; Riddle et al., 2020). Therefore, we tested whether EEG theta-band activity is primarily associated with the visual hemifield location of memory items via decoding analysis and examined resulting activation patterns mapped on EEG topography during significant decoding time.

We decoded the left-versus-right visual hemifield location of memory items. Upon target onset, we observed decoding performance significantly better than chance (peak Δ*p*(*decoding*) = 0.18 at 120 ms, lasting 280 ms; 1 cluster, *p* < 0.001; Figure 5e), similar to the results for decoding individual memory location (Figure 5b). The activation patterns mapped onto EEG topography during this period showed a clear binarization along the midline (Figure 5f), suggesting that decoding relied on differences in theta-band activity between the two hemispheres. During the delay period, significant decoding was observed regardless of whether the distractor was delivered (ND: peak Δ*p*(*decoding*) = 0.06 at 1060 ms relative to target onset, lasting 320 ms, 1 cluster, *p* < 0.001; WD: peak Δ*p*(*decoding*) = 0.06 at 1030 ms relative to target onset, lasting 300 ms, 1 cluster, *p* < 0.001; Figure 5g,i), consistent with our previous decoding results based on individual item locations. Significant decoding also began early when no distractor was shown (ND: from 800 ms to 1130 ms; WD: from 880 ms to 1180 ms). During this significant decoding time, time-resolved activation patterns in both trial conditions (Figure 5h,j) resembled those observed after target onset (Figure 5f), suggesting that improved item encoding at the anticipated distractor time may emerge through reactivation of the similar neural activity patterns seen during initial item encoding.

When we applied left-versus-right decoding on incorrectly performed trials, significant but less robust results were observed after either the anticipated distractor onset or actual distractor onset (ND: peak Δ*p*(*decoding*) = 0.05 at 870 ms, lasting 160 ms; 2 clusters, *p* < 0.001; WD: peak Δ*p*(*decoding*) < 0.001 at 1080 ms, lasting >110 ms; 1 cluster, *p* < 0.001; Supplementary Figure 4b). Compared to correctly performed trials (Figure 5g,i), decoding was weaker and its significant clusters were less contiguous, suggesting reduced information related to memory items during the delay period of incorrect trials.

We also decoded the upper-versus-lower visual hemifield locations of memory items and found significant decoding at target onset (Figure 5k), but for a shorter duration and with lower accuracy (peak Δ*p*(*decoding*) = 0.07 at 40 ms, lasting 150 ms; 1 cluster, *p* < 0.001) compared to left-versus-right decoding (peak Δ*p*(*decoding*) at 0.18 and lasted 280 ms). This decoding appeared to be driven more by the sensory response to the target stimulus than by memory encoding as decoding accuracy sharply peaked after target onset but immediately faded away. Consistent with this, we found no significant decoding after the anticipated or actual distractor onset (Figure 5m,o), and activation patterns were spurious over time (Figure 5n,p). These results indicate that EEG theta-band activity reflects more information about the encoding of left-versus-right visual hemifield locations compared to upper-versus-lower hemifield locations.

#### 5.3.3 Theta-band oscillations reflected reactivation of memory in anticipation of distractor

Because we found significant decoding of memory items both around target onset and during the anticipated distractor time, one natural question is whether the same neural activity patterns represent the memory item across these two distant periods. If so, this could be consistent with the same neural activity that originally encodes the target location being later “reactivated” in support of distractor resistance. Alternatively, it could be the case that representation of memory items during the delay relies on different neural activity patterns than those used for the initial encoding phase. To test whether the increased decoding performance after the anticipated time of the distractor stems from similar activity patterns during initial encoding phase at target onset, we applied a cross-temporal generalization analysis on the EEG theta-band activity.

During target onset period (0 - 500 ms), we found significant cross-temporal decoding between classifiers trained and tested within temporally neighboring time bins (clusters, *p* < 0.001; cluster-based permutation test with *α <* 0.05*, cluster α <* 0.05; N = 29 sessions combined across monkeys; Figure 6a,b), showing stability in the neural activity patterns representing the memory items over short time scales. Significant decoding was also observed when classifiers trained on data from the target onset period were evaluated on data from the anticipated (ND) or actual distractor (WD) onset period (700 - 1200 ms) (clusters, *p* < 0.001; Figure 6a,b). This showed that the classifiers were able to generalize learned activity patterns across different temporal phases of the task. When generalized within either the anticipated or actual distractor onset periods, similar results were obtained between conditions (clusters, *p* < 0.001; Figure 6a,b). However, the activity patterns were less generalized during the actual distractor onset period likely due to disrupting effect of the distractor (≥700 ms; Figure 6b), as shown from previous decoding results (Figure 5). When comparing between trial conditions (ND vs. WD), a significant difference was found when classifiers trained on the target onset period were generalized to the anticipated or actual distractor onset period (1 cluster, *p* = 0.024) (Figure 6c). No significant difference was observed in the reverse direction. This discrepancy may result from noise introduced by the distractor flash, which could reduce the information about the encoded memory items when training classifiers on those time bins.

**Figure 6:**
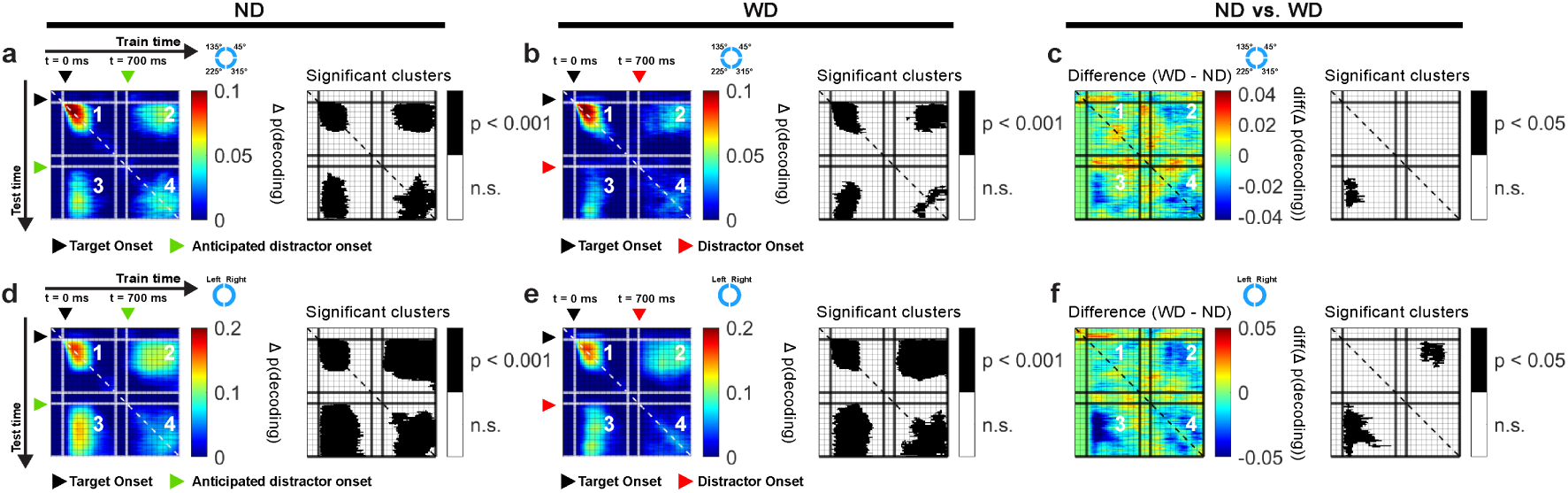
Cross-temporal generalization using EEG theta-band activity based on locations of items for ND (**a**) and WD (**b**) conditions, along with their difference (**c**). Similarly, cross-temporal generalization based on left and right visual hemifield location of items for ND (**d**) and WD (**e**) conditions with their difference (**f**). In each subplot, panel 1 generalizes within the post-target period (0 - 500 ms); panels 2 and 3 generalize between post-target period (0 - 500 ms) and post-anticipated-distractor or post-distractor period (700 - 1200 ms); panel 4 generalizes within post-anticipated-distractor or post-distractor period (700 - 1200 ms); corresponding significant clusters are shown in black-and-white maps. Black triangle: target onset; green triangle: anticipated distractor onset (ND); red triangle: distractor onset (WD). Color bars show decoding performance, in Δ p(decoding), above the chance level (0).

Since the left and right visual hemifield locations of memory items significantly influenced decoding performance (Figure 5e-j), we also performed a cross-temporal generalization analysis based on these locations (Figure 6d and e). Our findings largely mirrored those obtained from decoding individual memory item locations but showed larger significant clusters overall (clusters, *p* < 0.001). When comparing the ND and WD conditions, we found a significant difference when classifiers trained on the target onset period were generalized to either the anticipated or actual distractor onset periods (two clusters, *p* = 0.043, 0.002) (Figure 6f), indicating that the distractor interferes with memory reactivation.

In sum, cross-temporal generalization revealed that the EEG theta-band activity patterns were similar across distinct periods, consistent with “reactivating” memory items in the absence of a stimulus during the delay period of the task.

#### 5.3.4 Relative contributions of frontal and occipitoparietal theta-band activity in distractor anticipation during delay period

We aimed to disentangle the relative contributions of frontal and occipitoparietal EEG theta-band activity to distractor anticipation during working memory maintenance. To achieve this, we decoded the location of memory items using theta-band activity exclusively from either frontal or occipitoparietal EEG channels as classification features (groups shown in Figure 1b).

Decoding based on frontal theta-band activity showed a temporal profile similar to that obtained when using all EEG channels for classification (Figure 7a). At target onset, decoding was significant (peak Δ*p*(*decoding*) = 0.09 at 120 ms, lasting 260 ms; 1 cluster, *p* < 0.001), although slightly worse compared to using all EEG channels (peak Δ*p*(*decoding*) = 0.13; Figure 5b), suggesting that memory-related information is not just present in the frontal theta-band activity. After the anticipated or actual distractor onset, we found significant decoding that also displayed similar temporal profiles to the all-channel results but with decreased accuracy and a short time span (ND: peak Δ*p*(*decoding*) = 0.03 at 1000 ms, lasting 200 ms; 1 cluster, *p* < 0.001; WD: peak Δ*p*(*decoding*) = 0.02 at 1020 ms, lasting 160 ms; 1 cluster, *p* < 0.001).

**Figure 7:**
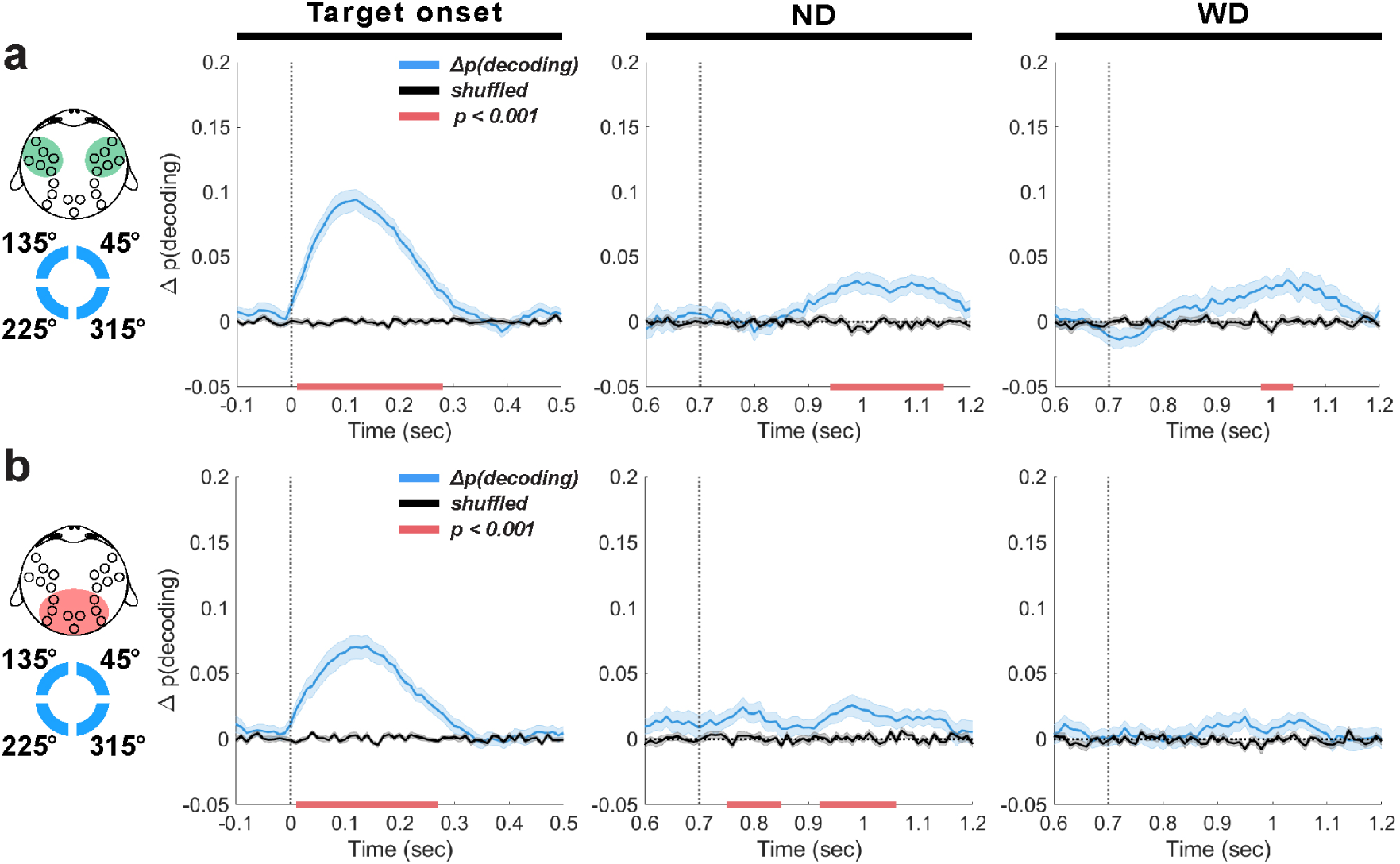
EEG theta-band decoding based on the frontal (**a**) or occipitoparietal EEG channel group (**b**). Decoding results (blue traces, ±1 SEM) are aligned to target onset (time = 0 ms), anticipated distractor onset (ND) and actual distractor onset (WD) (time = 700 ms) with their trial-shuffled results in black traces (±1 SEM). Horizontal red bars indicate significant decoding clusters (*p* < 0.001).

On the other hand, decoding using occipitoparietal theta-band activity resulted markedly different outcomes (Figure 7b). Shortly after the anticipated distractor onset, we observed weak but significant decoding (ND: peak Δ*p*(*decoding*) = 0.02 at 780 ms, lasting 90 ms; 1 cluster, *p* < 0.001), followed by a similar significant decoding 220 ms later (peak Δ*p*(*decoding*) = 0.02 at 930 ms, lasting 120 ms; 1 cluster, *p* < 0.001). While these effects were weak, they imply that memory-related information emerges earlier in the occipitoparietal area compared to the frontal area. This decoding never reached significance when the distractor was actually delivered. This differs from the frontal theta-band decoding that showed robust performance even with distractor presentation. These findings reflect the disruptiveness of the distractor on memory representations in the occipitoparietal area, consistent with previous findings showing reduced memory representation in the visual cortex following distractor presentation (Bettencourt and Xu, 2016; Xu, 2017).

In summary, we found that theta-band activity in the frontal area conveyed more information about memory items than in the occipitoparietal area. The temporal profile of decoding using theta-band activity from frontal EEG group resembled that of decoding using theta-band activity collected from all EEG electrodes. Although weak, decoding from the occipitoparietal theta-band activity in the absence of stimuli may suggest the emergence of memory-related signals during distractor anticipation. This weak decoding may be attributed to the simple nature of our target visual stimulus (i.e., a circle) that may not require significant involvement of the occipitoparietal area. These results suggest while both frontal and occipitoparietal areas involve in enhanced representation of memory items during distractor anticipation, a significantly greater contribution is made from the frontal area.

### 5.4 Prefrontal LFP theta-band activity in working memory maintenance and distractor anticipation

Because the LFP recorded from LPFC also showed significant theta-band activity, we hypothesized that the LPFC theta-band activity is also associated with individual memory items during the task. We also tested whether the task-irrelevant checkerboard modulated activity in a way that allowed inferences about the memory item. Such a memory-dependent modulation would be consistent with latent representation of memory items (Stokes, 2015). We also hypothesized that if the LPFC involves in enhancement of memory encoding during distractor anticipation, we would observe increased decoding of items. To test these, we decoded memory items based on their (1) individual locations and (2) distributions across visual hemifields using the LFP theta-band activity recorded from the LPFC.

#### 5.4.1 LPFC theta-band activity reflects the encoding of item in response to a distractor

We tested whether LFP theta-band activity of the LPFC is associated with encoding of memory items by decoding based on their individual locations. At target onset, we found accurate decoding well above chance in both monkeys (monkey R: peak Δ*p*(*decoding*) = 0.08 at 190 ms, lasting 380 ms, 1 cluster, *p* = 0.002; monkey W: peak Δ*p*(*decoding*) = 0.11 at 100 ms, lasting 300 ms, 1 cluster, *p* < 0.001; Figure 8a,b). During the delay period, significant decoding was also detected in response to the distractor (monkey R: peak Δ*p*(*decoding*) = 0.09, onset 800 ms, lasting 130 ms, 1 cluster, *p* = 0.002; monkey W: peak Δ*p*(*decoding*) = 0.05 at 800 ms, lasting 270 ms; 1 cluster, *p* < 0.001; Figure 8a,b; WD). However, when the distractor was not shown, decoding performance generally did not reach significance in either monkey, except for a short duration (approximately 50 ms) in monkey R a few hundred milliseconds after the anticipated distractor onset (peak Δ*p*(*decoding*) = 0.04 at 950 ms; 1 cluster, *p* = 0.003; Figure 8a; ND). Overall results suggest that improved memory encoding in anticipation of a distractor was not reflected in LPFC theta oscillations, which contrasted to our EEG theta-band results that showed significant decoding around the anticipated distractor time (Figure 5c,d). Additionally, our finding that the distractor not associated with the target visual stimulus improved the encoding of items implicates that the LFP theta-band response to the distractor reveals memory items encoded by underlying LPFC neurons, consistent with prior research that demonstrated that the LPFC neurons’ firing rate reflect memory-dependent response to a task-irrelevant visual stimulus, showcasing its capacity to reveal the latently represented memory in the LPFC (Stokes et al., 2013).

**Figure 8:**
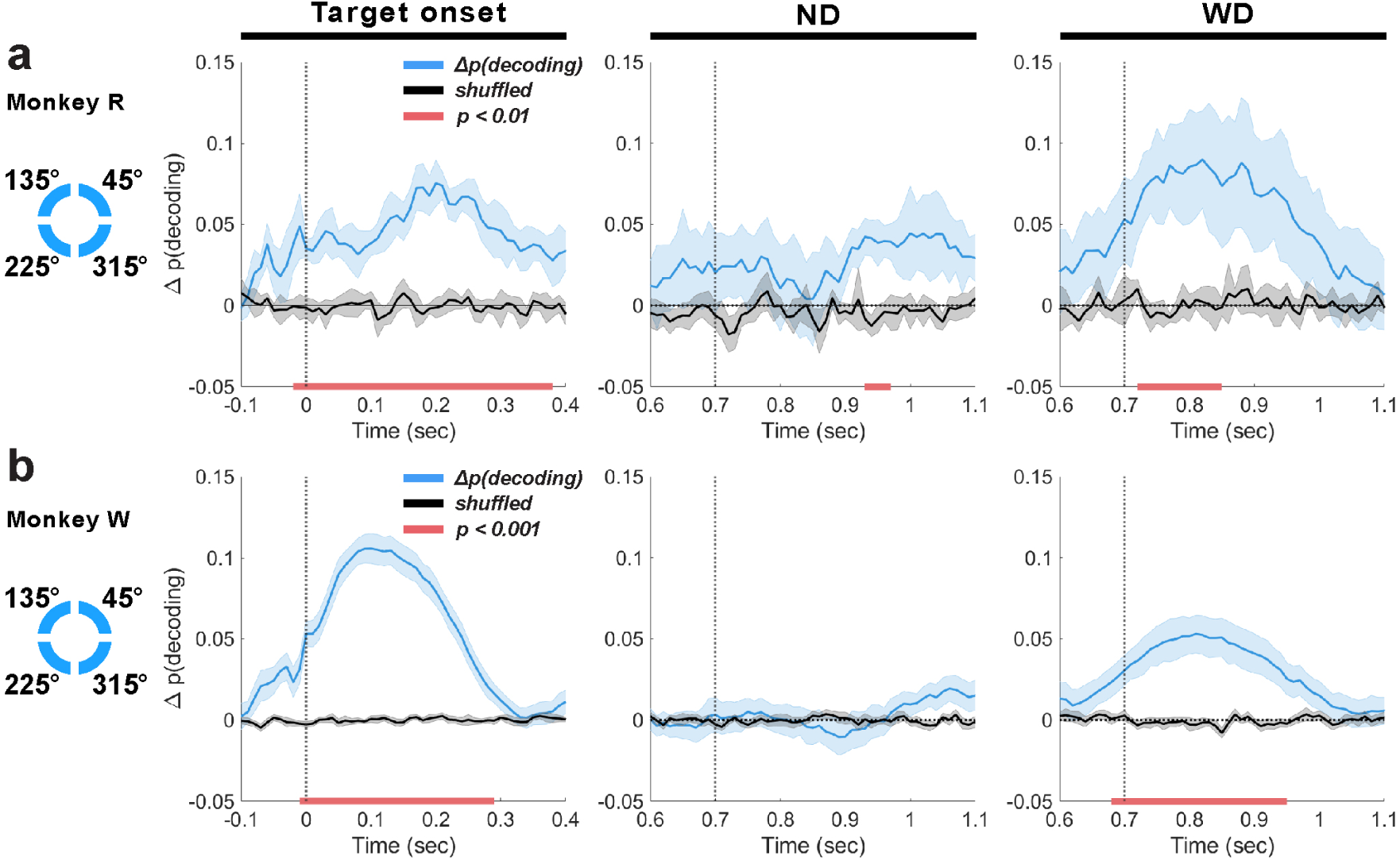
Decoding based on the location of individual memory items using LFP theta-band activity (4 - 8 Hz) for monkey R (**a**) and W (**b**), aligned to target onset (t = 0 ms), anticipated distractor onset (ND; t = 700 ms), and actual distractor onset (WD; t = 700 ms). Mean decoding performance (blue traces, ±1 SEM) and trial-shuffled performance (black traces, ±1 SEM) are shown, with significant clusters (*p* < 0.01 or *p* < 0.001 depending on monkey) indicated by horizontal red bars.

To determine whether the significant decoding from the LPFC theta-band activity actually reflected memory encoding rather than mere sensory processing, we attempted to decode memory items from LFPs during incorrectly performed trials using classifiers trained on data from correctly performed trials. If the theta-band activity simply reflected the feed-forwarding processing of the target visual stimulus, we would observe increased decoding at target onset whether the monkeys performed the task correctly or not. However, when the monkeys performed the task incorrectly, we did not find significant, contiguous decoding at target onset, indicating that the decoded information reflects the memory being encoded after a brief stimulus presentation (Supplementary Figure 5). Furthermore, unlike during correctly performed trials, distractor presentation did not improve decoding performance, remaining at chance.

This analysis supports that the improved decoding observed during target and distractor presentation in the correctly performed trials reflected encoding of memory items.

To test which location properties of items were primarily reflected by the theta-band activity, we performed decoding analysis for all pairwise combinations of possible item locations (_4_*C*_2_ = 6 pairs). For all combinations, no significant cluster was found immediately following the anticipated distractor onset (time ≥700 ms) (Supplementary Figure 6a,b; ND). But at the later time period (time ≥900 ms), significant decoding was observed in monkey R but without a clear pattern (clusters; 45*^◦^* vs. 135*^◦^*: *p* = 0.008, 45*^◦^* vs. 225*^◦^*: *p* = 0.04, 45*^◦^* vs. 315*^◦^*: *p* = 0.002, 135*^◦^* vs. 315*^◦^*: *p* = 0.03, 225*^◦^* vs. 315*^◦^*: *p* = 0.01) and in monkey W (45*^◦^* vs. 315*^◦^*: *p* = 0.004). Although subtle, this increased decoding during the anticipated distractor time appeared similar to what we observed from the EEG decoding during this time period (Figure 5g). In monkey W, a transient decoding was observed in the later time (≥1000) only when comparing 45*^◦^* vs. 315*^◦^* (*p* = 0.02).

When the distractor was actually presented, significant decoding was achieved in both monkeys especially when the items were decoded based on their left or right visual hemifield location (45*^◦^* vs. 135*^◦^*: monkey R, *p* = 0.008, monkey W, *p* = 0.001; 45*^◦^* vs. 225*^◦^*: monkey R, *p* = 0.002, monkey W, *p* = 0.003, 135*^◦^* vs. 315*^◦^*: monkey R, *p* = 0.02, monkey W, *p* < 0.001, and 225*^◦^* vs. 315*^◦^*): monkey R, *p* = 0.002, monkey W, *p* = 0.003) (Supplementary Figure 6a,b; WD). And, in general, no significant effect was found when decoding memory items that were shown in the same visual hemifield (i.e., 45*^◦^* vs. 315*^◦^* and 135*^◦^* vs. 225*^◦^*), except in monkey W, where a significant effect (1 cluster, *p* < 0.001) was observed when decoding items within the hemifield contralateral to the microelectrode implant site (i.e., 135*^◦^* vs. 225*^◦^*).

Overall, These results indicate that LPFC LFP theta-band activity also predominantly reflects left or right visual hemifield locations of memory items, extending our findings from EEG analysis (Figure 5).

#### 5.4.2 Prefrontal theta-band oscillations did not show reactivation of memory during distractor response

We tested whether the theta-band activity patterns between the target and distractor time periods were similar. We applied cross-temporal generalization based on individual item locations (Figure 9). In short, we did not find a similar activity pattern between the two distinct time periods. When the distractor was shown (time = 700 ms), we found significant decoding clusters, generalized within neighboring time bins (monkey R: 1 cluster, *p* = 0.009; monkey W: 1 cluster, *p* < 0.001; Figure 9b,e; panel 4). These results suggested that different mixtures of LPFC neurons encode items (i.e., dynamic coding) during the delay period compared to the post-target period, as suggested by previous studies (Stokes et al., 2013). In the ND condition, we observed a difference between monkeys where a significant decoding cluster (*p* = 0.009) was detected in monkey R (Figure 9a; panel 4) but not in monkey W (Figure 9d; panel 4). Also, when compared between trial conditions, no difference was observed in monkey R (Figure 9c) but in monkey W (1 cluster, *p* = 0.02) (Figure 9f).

**Figure 9:**
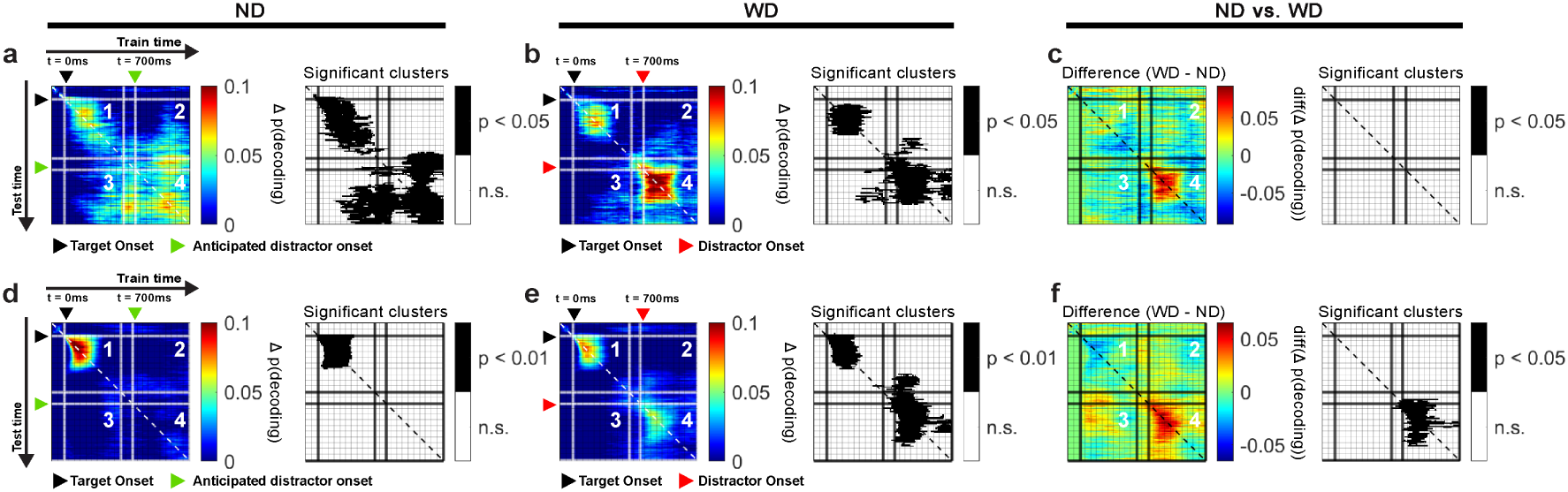
Cross-temporal generalization of LPFC LFP theta-band activity for monkeys R and W, based on item locations in ND (**a** and **d**) and WD (**b** and **e**) conditions, with their differences (ND vs. WD) (**c** and **f**). Panel 1 generalizes within the post-target period (0 ms - 500 ms). Panels 2 and 3 generalize between the post-target (0 ms - 500 ms) and post-anticipated-distractor/post-distractor periods (700 ms - 1200 ms). Panel 4 generalizes within the post-anticipated-distractor/post-distractor period (700 ms - 1200 ms). Significant decoding clusters shown in black-and-white maps. Black triangle: target onset; red triangle: distractor onset (in WD); green triangle: anticipated distractor onset (in ND). Color bars show decoding performance, Δ p(decoding), above the chance level; values below chance are not shown.

We also attempted to test whether the LPFC LFP beta-band activity possibly reflects memory items during the delay period and generalize across time. We found no evidence of this, as significant decoding appeared only at target onset (Supplementary Figure 7).

To summarize, we observed increased encoding of memory items specifically in the LPFC theta-band activity in response to target and distractor stimuli. This implicates that the underlying memory items encoded by LPFC neurons becomes revealed when activated by an external impulse stimulus like our checkerboard visual stimulus, consistent with prior research demonstrating that a visual stimulus impulse that is irrelevant to working memory tasks elicits neural responses that are dependent on encoded memory (Stokes et al., 2013; Stokes, 2015). Moreover, unlike our EEG findings (Figure 6), we found no evidence of memory reactivation during distractor anticipation.

### 5.5 Functional Connectivity between the LPFC and the occipitoparietal cortex: Strong intrahemispheric connections

Previous studies have reported that local rhythmic activity can drive interareal synchronization between the frontal cortex and other brain areas (Fries, 2005; Bastos et al., 2015). In the context of working memory, interactions between the LPFC and occipitoparietal brain areas are suggested to be crucial for maintenance and control of information (Salazar et al., 2012; Antzoulatos and Miller, 2016; Jacob et al., 2018). Since both LFP and EEG originate from the same underlying biophysical processes but are sampled at different spatial scales (Buzsáki et al., 2012), we wanted to examine how local activity in the LPFC influences other brain areas during distractor anticipation. To test this, we simultaneously recorded LFP from the LPFC and EEG from the scalp, and estimated coherence and Granger causality index (GCI) between these signals.

#### 5.5.1 Beta-band activity facilitates intrahemispheric, long-range functional connectivity between the LPFC and the occipitoparietal cortex

While our decoding analysis showed that theta-band activity across spatial scales was most strongly associated with working memory maintenance and distractor anticipation (Figures 5 and 8), our coherence analysis instead showed significance predominantly in the beta band (13 - 30 Hz) (Figure 10).

**Figure 10:**
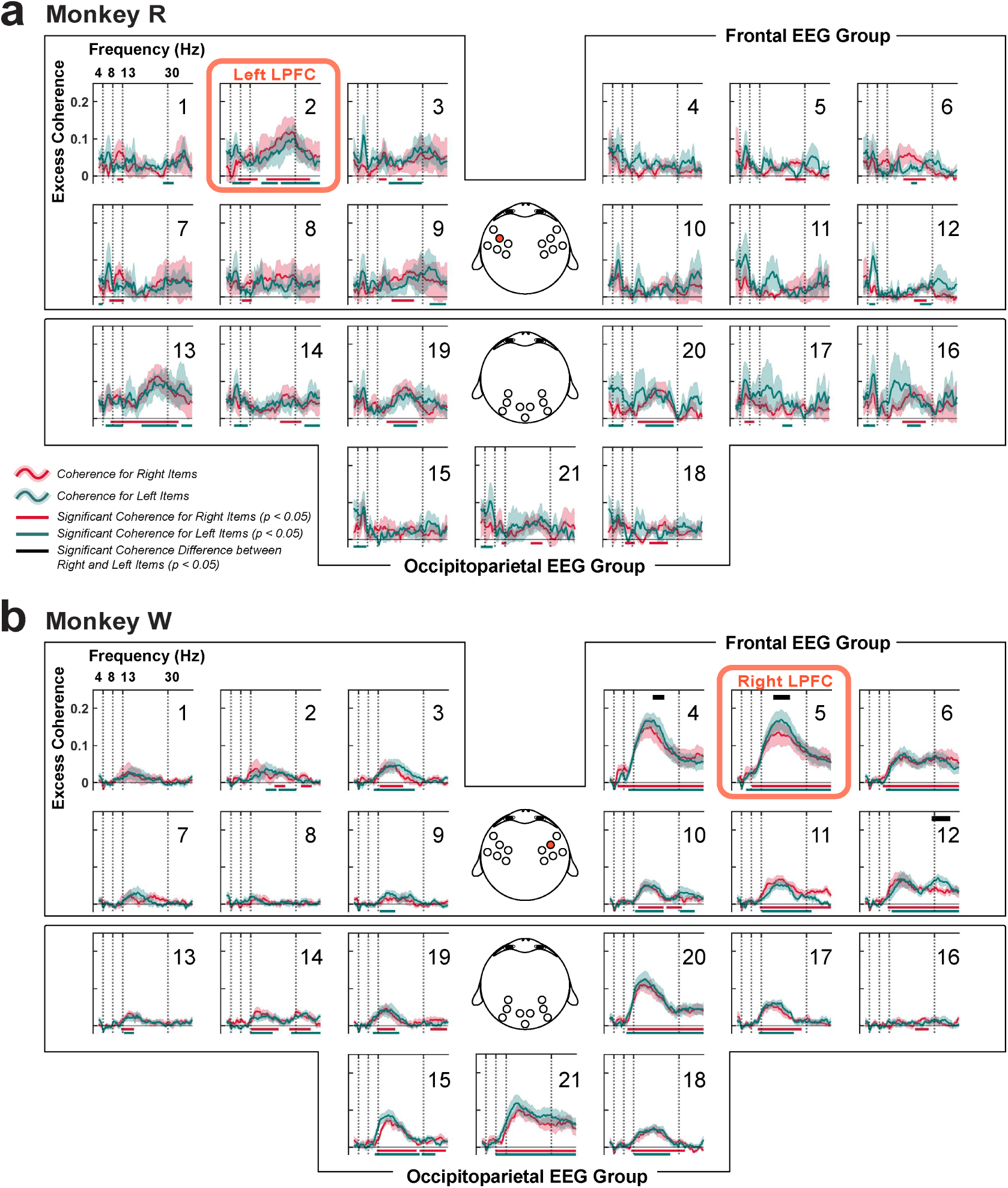
Coherence between the LPFC LFP and scalp EEG, organized into frontal and occipitoparietal EEG channel groups. (**a**) Monkey R had the microelectrode array implanted in the left LPFC near EEG channel 2 (red box; highlighted on the topography in the middle). (**b**) Same for monkey W that had the array implanted in the right LPFC near EEG channel 5 (red box; also highlighted). Red traces (±1 SEM) represent coherence while remembering items presented in the right visual hemifield, and green traces (±1 SEM) represent coherence when remembering items shown in the left visual hemifield. Corresponding significance is shown in their respective colors at the bottom (clusters, *p* < 0.05). Significant differences between item conditions are displayed in the black horizontal bars at the top (clusters, *p* < 0.05).

Within the frontal EEG group, we found significant beta-band coherence between the LPFC LFP and the EEGs collected near the LFP recording sites (monkey R: EEG channel 2, *p* < 0.05, N = 6 sessions; monkey W: EEG channels 4, 5, and 6 with *p* < 0.05. N = 7 sessions; cluster-based one-sample, one-tailed permutation test with *α <* 0.05*, cluster α <* 0.05), similar to a previous study using a visual change-detection task (Snyder et al., 2018). We compared between item conditions by computing coherence after grouping items based on their left or right visual hemifield location. Only in monkey W, we found significant differences between the conditions in the frontal EEG group, specifically at those EEG electrodes around the LFP array implant site (channels 4, 5, and 12 with *p* = 0.02, 0.04, and 0.02, respectively; N = 7 sessions; cluster-based two-sample, two-tailed permutation test with *α <* 0.05*, cluster α <* 0.05; Figure 10b). As left items were items shown contralateral to the LPFC implant site in monkey W, this suggested that the higher coherence is observed when remembered contralateral items. However, such was not observed in monkey R.

In addition, significant excess beta-band coherence was observed broadly across the occipitoparietal brain area (monkey R: EEG channels 13, 14, 16, 19, 20, and 21, clusters, *p* < 0.05, N = 6 sessions; monkey W: EEG channels 13, 14, 15, 17, 18, 19, 20, and 21, clusters, *p* < 0.05). But no significant difference between item conditios were found. For both monkeys, these coherence values tended to be higher on the hemisphere ipsilateral to the LPFC implant site. This enhanced coherence between the LPFC and occipitoparietal area may play a crucial role in distractor anticipation, possibly to support inter-areal communication to remember the maintained representation of working memory items.

To determine the directionality of interareal communication, we computed GCI between the LPFC LFP and scalp EEG (Figure 11). Overall, GCI was less noisy in monkey W compared to monkey R. In both monkeys, we observed significant GCI, also predominantly in the beta band, consistent with coherence analysis. GCI from the LPFC LFP to the overlying scalp was greater than GCI in the other direction (monkey R, N = 6 sessions, EEG channel 2, 1 cluster, *p* = 0.04; monkey W, N = 7 sessions, EEG channel 5, 1 cluster, *p* = 0.03; cluster-based one-sample, one-tailed permutation test with *α <* 0.05*, cluster α <* 0.05). Additionally, in monkey W, several EEG channels overlying near the LPFC LFP recording site showed significant beta-band GCI from the LFP (clusters, channels 6, 11, and 12 each with *p* = 0.02).

**Figure 11:**
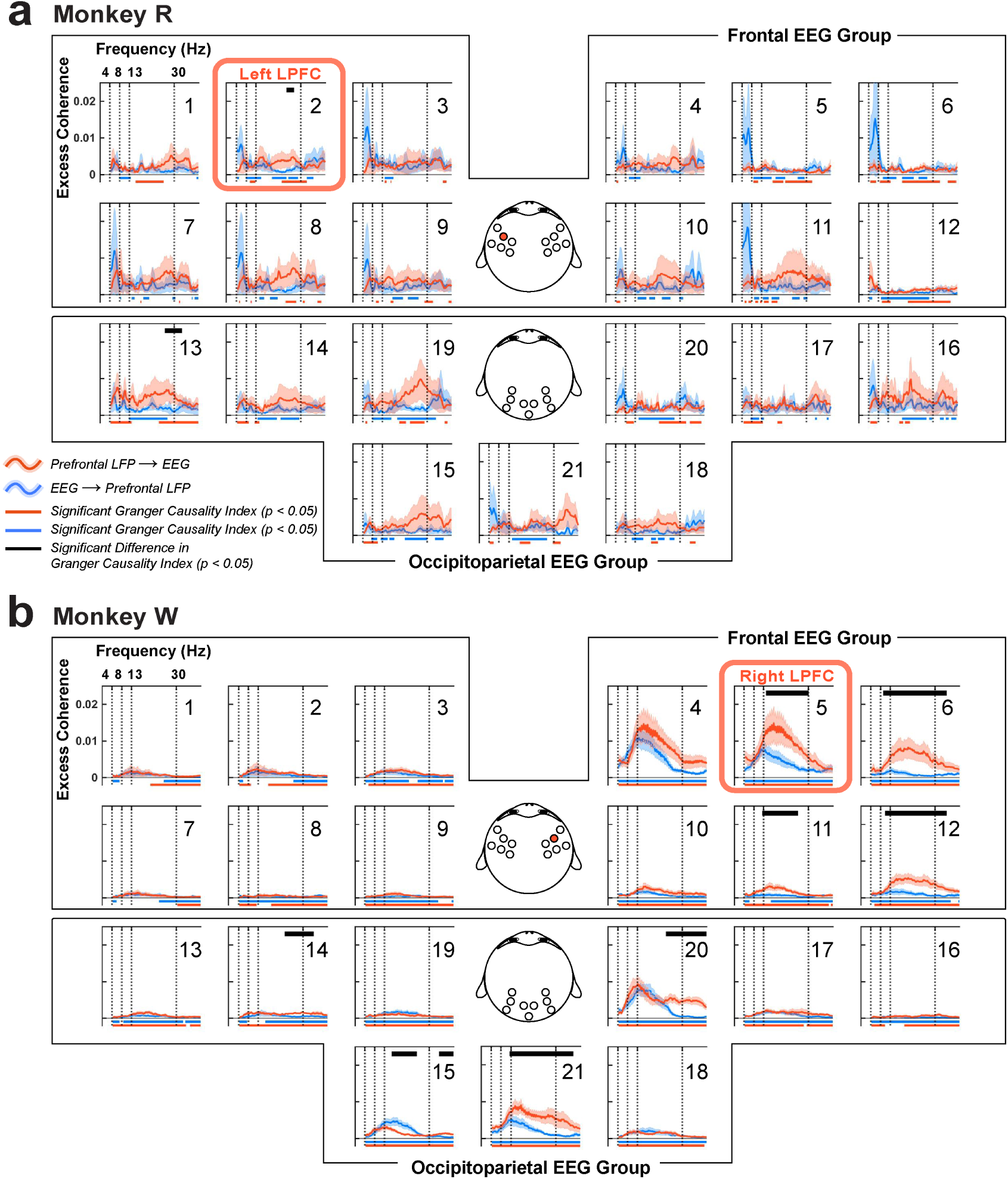
Granger causality indices (GCI) between the LPFC LFP and scalp EEG, organized into frontal and occipitoparietal EEG channel groups. (**a**) Monkey R received a microelectrode array implant in the left LPFC near EEG channel 2 (red box and also highlighted on the EEG topography). (**b**) For monkey W, the implant site was in the right LPFC, near EEG channel 5 (red box and highlighted in the corresponding EEG topography). Red traces (±1 SEM) represent GCI from LPFC LFP to EEG, while blue traces (±1 SEM) indicate the reverse direction, from EEG to LPFC LFP. Significant GCI are marked with horizontal bars at the bottom in their respective colors (clusters, *p* < 0.05). Significant differences between the GCIs are indicated by black horizontal bars at the top (clusters, *p* < 0.05).

We also found significant beta-band GCI from the LPFC LFP to the EEG overlying the occipitoparietal area (monkey R, N = 6 sessions, EEG channel 13, 1 cluster, *p* = 0.03; monkey W, N = 7 sessions, clusters, channels 14, 15, 20, and 21 with *p* = 0.03, 0.008, 0.04, and 0.02, respectively) (Figure 11a,b; occipitoparietal EEG group). In monkey R, EEG channel 19 showed a trend toward a significant top-down GCI but did not reach significance (however, this LFP-EEG pair was significantly coherent; Figure 10a). Our findings align with previous studies (Bastos et al., 2015), highlighting the role of beta-band oscillations in top-down feedback to the occipitoparietal area. In monkey W, we saw one LFP-EEG pair with a significant bottom-up GCI compared to its top-down GCI in the beta band (Figure 11b; EEG channel 15, clusters, *p* = 0.04); however the source of this bottom-up GCI was unclear and beyond the scope of this paper. Across all monkeys, theta and alpha bands showed no differences in GCIs across directions. Importantly, these directed frequency-domain interactions were mainly observed in the brain areas ipsilateral to the LPFC LFP recording site, showing that the LPFC predominantly exert top-down signaling in the ipsilateral occipitoparietal brain areas.

#### 5.5.2 Interareal beta-band coherence briefly fluctuates in anticipation of a distractor during the delay period

To better understand the spatial distribution of the beta-band coherence, we constructed LFP-EEG coherence topography by averaging the largest coherence (Coherence_max_) within the beta band across recording sessions for each LFP-EEG pair for each monkey (Figure 12a,b). We used the maximum value within the beta band because the number of peaks and troughs appeared varying at different frequencies across monkeys, as shown in our power spectra examples (Figure 4a,b). Taking the maximum ensured that we capture the most prominent oscillatory feature in the desired frequency band, which could be more representative of the neural activity’s peak influence compared to a mean, which might obscure these key fluctuations.

**Figure 12:**
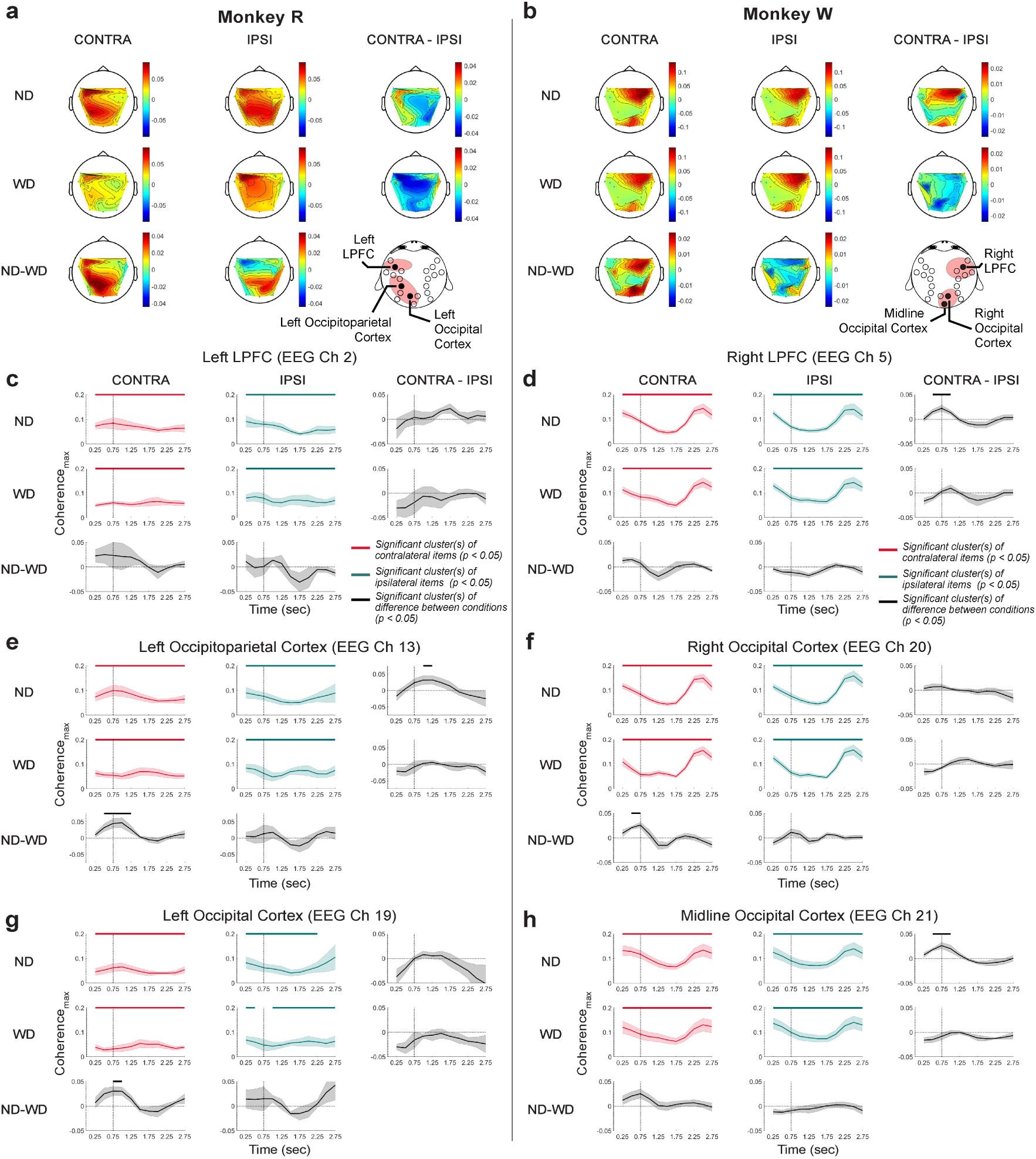
Beta-band coherence topography and time-resolved coherence between the LPFC LFP and scalp EEG. The vertical line at 700 ms marks the time of either anticipated or actual distractor onset. The monkeys R (**a**) and W (**b**) topographies for each item (CONTRA and IPSI) and trial condition (ND and WD) with their differences across conditions (CONTRA-IPSI and ND-WD). EEG channels that showed enhanced coherence with the LFP are marked on the montage at the bottom left corner. (**c**,**e**,**g**) Time-resolved LFP-EEG coherence_max_ (±1 SEM) for monkey R across trial and item conditions. (**d**,**f**,**h**) Same but for monkey W. CONTRA (in red traces) refers to trials where monkeys remembered items displayed in the visual hemifield contralateral to the LPFC implant site, while IPSI (in green traces) refers to trials with items displayed in the visual hemifield ipsilateral to the implant site. Differences across conditions in black traces (±1 SEM). Significance indicated by horizontal bars (cluster-based permutation test result with *α* = 0.05 and *cluster α* = 0.05) at the top of each plot with their respective colors.

We compared the coherence topography across trial conditions (ND vs. WD) and relative item locations (contralateral vs. ipsilateral to the LPFC array implant site), and found both monkeys showed subtly stronger intrahemispheric coherence when remembering items displayed on the contralateral visual hemifield (CONTRA) than when remembering items displayed on the ipsilateral side (IPSI).(Figure 12a,b; ND, CONTRA-IPSI). When the distractor was displayed, no difference was detected (WD, CONTRA-IPSI), likely occluded by the strong effect of the distractor in both time and frequency domains, as shown in our previous analysis (Supplementary Figure 2).

We next wished to test the time course with which LFP-EEG coherence evolved over time in a small window over time (i.e., 500-ms window with a 250-ms overlapping step). We restricted this analysis to the LFP-EEG pairs that most exhibited significant difference in coherence between CONTRA and IPSI conditions in the beta-band coherence topographies. To be specific, in monkey R, we calculated LFP-EEG coherence with EEG channels 2, 13, and 19, which approximately overlie the left LPFC, left occipitoparietal cortex, and left occipital cortex, respectively. For monkey W, we used channels 5, 20, and 21, corresponding to the right LPFC, right occipital cortex, and midline occipital cortex, respectively.

When monkey R performed the task correctly, coherence between the LFP and EEG over the left LPFC (channel 2) stayed significant above chance throughout the delay period of the task (clusters, *p* < 0.01; a cluster-based one-sample, one-tailed test with *α* < 0.05, *cluster α* < 0.05). However, no significant differences emerged between conditions (Figure 12c), despite a subtle tendency to increase when remembering contralateral versus ipsilateral items in the later time of the delay period (1700 ms; ND, CONTRA-IPSI). Most importantly, we found a significant, transiently increased coherence with the EEG over the left occipitoparietal cortex (channel 13) around the anticipated distract time (1000 ms) in CONTRA items compared to IPSI items (1 cluster, *p* = 0.02, N = 6 sessions; a cluster-based two-sample, two-tailed permutation test with *α* < 0.05, *cluster α* < 0.05; Figure 12e; ND, CONTRA-ISPI). Distractor presentation briefly reduced interareal coherence during CONTRA item maintenance (1 cluster, *p* = 0.03; ND-WD, CONTRA) but not for IPSI items, suggesting that the coherence between the two brain areas are modulated specifically when memory item shown in the contralateral visual hemifield is being retained.

In monkey W, similar results were obtained. Across trial conditions, coherence between EEG overlying the right LPFC site (channel 5) and the LPFC LFP was significant through out the task (clusters, *p* < 0.001, N = 7 sessions), but showed a different temporal profile compared to monkey R. Strength of coherence progressively decreased until the end of the delay period (<2000 ms) and abruptly increased during saccade (>2000 ms) (Figure 12d). Importantly, stronger coherence was detected during the anticipated distractor time (around 750 ms) when remembering contralateral versus ipsilateral items (1 cluster, *p* = 0.003; ND, CONTRA-IPSI). Similarly, such transient coherence increase was also observed around the anticipated distractor time between EEG over the midline occipital area and the LFP (channel 21; around 750 ms, 1 cluster, *p* = 0.01; Figure 12h; ND, CONTRA-IPSI).

When the monkeys failed to perform the task correctly, transient coherence increase around the anticipated distractor time was not present in all representative LFP-EEG pairs (Supplementary Figure 8; ND, CONTRA-IPSI for every LFP-EEG pairs). This suggests that enhanced communication between the LPFC and occipitoparietal area at the time of the anticipated distractor may be crucial to successfully carry out the task.

We also investigated the behavior relevance of the transiently increased coherence between the LPFC and occipitoparietal areas by correlating beta-band coherence with trial-level behavioral error estimates; saccadic reaction time and memory recall error. However, no significant correlations were found (Supplementary Figure 9). In addition, we analyzed the relationship between these error estimates and mean LFP band power, but no consistent or coherent effects were detected across monkeys (Supplementary Figure 10). This suggests that the distractor had negligible influence on systematic biases in memory signals in the brain.

To summarize, we found enhanced interareal communication between the LPFC LFP and the EEG over the occipitoparietal cortex, primarily within the same hemisphere. This communication was particularly notable around the anticipated distractor time, especially when remembering the items displayed contralateral to the LPFC location. While this increase did not correlate with trial-wise behavioral errors, it seemed pertinent for successful carrying out the task. These results suggest the fronto-occipitoparietal coherence is modulated task-dependently during distractor anticipation.

## 6 Discussion

The present study investigated how brain oscillations in monkeys across spatial scales –prefrontal LFP and scalp EEG –are associated with distractor anticipation during working memory maintenance. Theta-band oscillations were closely associated with the item being remembered during the delay period of the MGS task. Large-scale theta-band oscillations (EEG) also responded when anticipating distraction –consistent with memory reactivation –a pattern not observed in smaller-scale theta-band oscillations (LFP) within the LPFC via decoding and cross-temporal generalization analyses. EEG theta-band activity shortly after (<100 ms), compared to before, the anticipated distractor onset more effectively encoded items for a short period (<500 ms), with only subtle disruption during the presentation of a distractor. In contrast, LPFC LFP theta-band oscillations showed increased item encoding only briefly after distractor onset (<100 ms), revealing the memory item latently represented by underlying LPFC neurons (e.g. Stokes, 2015). Across spatial scales, items were predominantly encoded by their left or right visual hemifield locations, reflecting the brain’s organization of spatial information during working memory. In addition, we analyzed LPFC LFP and scalp EEG together and found enhanced interareal communication between the ipsilateral LPFC and occipitoparietal areas, specifically in the beta band, during working memory maintenance, as indicated by our coherence and Granger causality analyses. Intriguingly, this intra-hemispheric communication transiently strengthened around the anticipation of distractor onset, reflecting a possible preparatory mechanism towards the anticipated distractor. Together, these findings show that distractor anticipation affects neural oscillatory activity during working memory maintenance, with distinct effects across spatial scales and frequency bands, potentially protecting working memory content from forthcoming distraction.

Humans and monkeys, despite evolutionary divergence, exhibit similar oscillatory characteristics due to structural and functional homologies (Petrides and Pandya, 2002; Neubert et al., 2014; Buzsáki et al., 2013). Comparative studies between humans and monkeys showed similar oscillatory behaviors during working memory (Reinhart et al., 2012), bridging gaps between human and monkey electrophysiology. Distractor anticipation during working memory maintenance suppresses alpha-band oscillations in humans, suggesting their role in mitigating distractor-induced interference (Foxe and Snyder, 2011; Bonnefond and Jensen, 2012; Gutteling et al., 2022; Magosso and Borra, 2024). This human alpha is also associated with visuospatial attention (Snyder and Foxe, 2010; Worden et al., 2000; Rihs et al., 2007) and working memory (Foster et al., 2016; Bae and Luck, 2018), providing neural signatures to track either the attended or memorized locations. Therefore, in monkeys, we initially expected to see task-related activity predominantly in the alpha band during our spatial working memory task. However, we did not detect predominant alpha-band oscillations (Figure 4; Supplementary Figure 2), unlike some prior works in monkey neurophysiological studies (Buschman et al., 2012; Reinhart et al., 2012; Lara and Wallis, 2014; Snyder et al., 2015; Holmes et al., 2018; but see Wimmer et al., 2016a). One explanation is that the brain areas or cortical depth (i.e., superficial layer) we recorded from simply do not show dominant alpha band activity (Mendoza-Halliday et al., 2024a), thereby PCA could not capture the activity. Another possibility is that the monkey homolog of the human alpha band ranges across wider frequencies, from 8 Hz to 16 Hz or even higher, overlapping with the beta band that ranges between 13 Hz and 30 Hz. While some activity was observed in the range but no clear-cut boundaries were found to confidently isolate the alpha from the beta in both LFP and EEG data (Figure 4c,j; 12 - 40Hz). In this study, we instead found significant task-related oscillations in the theta and beta bands, just outside the traditional alpha band range.

Studies with human participants have previously shown strong associations between the theta-band activity and working memory, such as increased theta-band power in proportion to increased memory load (e.g., number of items being remembered) (Gevins et al., 1997; Raghavachari et al., 2001; Jensen and Tesche, 2002; Cohen and Donner, 2013) and modulated working memory performance with theta-band-based brain stimulation (improvement studies: Polanía et al., 2012; Hoy et al., 2016; Albouy et al., 2017; Violante et al., 2017; Riddle et al., 2020; disruption studies: Lee and D’Esposito, 2012; no effect study: Biel et al., 2022). In monkeys, we as well observed that the scalp EEG theta-band oscillations were pronounced during the MGS task and, interestingly, exhibited transient memory-dependent activities after the anticipated distractor time that were only briefly disrupted by the distractor (Figure 5). Fuentemilla and colleagues (2010) previously demonstrated that the theta-band oscillations from human magnetoencephalogram (MEG) reflect a periodic replay of visual information being remembered during a delayed match-to-sample working memory task (Fuentemilla et al., 2010). What we observed from monkeys is replay-like (that we called memory reactivation in this paper) as demonstrated by our cross-temporal generalization results showing that the neural activity patterns during the post-target period resemble the patterns during the post-anticipated-distractor period (Figure 6). Furthermore, decoding during this period was not effective when the monkeys failed to successfully carryout the task (Supplementary Figure 4). This may suggest that the memory replay is related to successful execution of working memory task during possible distraction. And during this memory reactivation, we saw significant contributions from the frontal theta-band activity (Figure 7). Weaker contributions from the occipitoparietal theta-band activity could be attributed to the simplicity of our target visual stimulus –a circle –which may not demand substantial involvement of the modality-specific brain areas, here the visual cortex, for information maintenance as implicated by the sensory recruitment hypothesis.

In the LPFC, we did not observe memory reactivation in theta-band activity. This may be due to lack of memory-dependent theta-band activity in the LPFC (e.g., areas 8A and 9/46) during the delay period of working memory tasks (Wimmer et al., 2016a; Holmes et al., 2018). But, when a distractor was delivered, we could decode memory items from the theta-band response (Figure 8). Since the distractor was constant in appearance and timing, and, importantly, lacked spatial relevance to memory items, the decoded information was likely associated with the latent memory representation encoded by LPFC neurons, consistent with prior studies (Stokes et al., 2013; Wolff et al., 2017; Ten Oever et al., 2020). This was further supported by the lack of decoding in incorrect trials at both target and distractor onsets (Supplementary Figure 5). In addition, our cross-temporal generalization results showed the representation of memory items evolved over time (Figure 9), aligning with the “dynamic coding” framework (Zaksas and Pasternak, 2006; Stokes et al., 2013; Stokes, 2015), in which different mixtures of neurons encode throughout memory maintenance from the changes in their underlying functional connectivity and tuning properties (Duncan, 2001). This may explain the absence of memory reactivation after the anticipated distractor onset, as different neurons mediate initial encoding and maintenance, leading to distinct theta-band activity over time.

In the LPFC, we also observed anticipatory suppression particularly in the theta band, which endured until the anticipated distractor time and was followed by a rapid recovery to a pre-distractor level (Supplementary Figure 3). This is similar to a previous study showing that LPFC neurons suppress their spiking activity in anticipation of a distractor, with the strongest suppression occurring when the distractor was displayed farthest from remembered target location (Suzuki and Gottlieb, 2013). Possibly, anticipatory processes may shape LPFC activity distractor-dependently, akin to human non-invasive studies that showed that the salience of distractor modulates brain oscillations at different intensity (Bonnefond and Jensen, 2012; Magosso and Borra, 2024), but not memory-dependently. The recovery of suppressed theta-band oscillations after the anticipated distractor onset may spontaneously enhance theta-band-based coupling (Liebe et al., 2012; Fuentemilla et al., 2010) or synchronization (Sarnthein et al., 1998) widely across the hemisphere, leading to memory replay that we observed in the EEG theta-band oscillations. However, during this recovery period, interareal theta-band coherence with respect to LPFC (i.e., LFP-EEG coherence) was weak and not distinct between the LPFC and EEG (Figure 10). Instead, we found task-related coherence specifically from the beta-band oscillations between the LPFC and occipitoparietal area.

We found significant beta-band coherence between the LPFC LFP and scalp EEG across monkeys (Figure 10), with Granger causal influence predominantly from the LPFC to ipsilateral occipitoparietal areas (Figure 11). The pattern of coherence between the LPFC and EEG subtly differed across monkeys, likely reflecting hemispheric asymmetry in connectivity and functions (Falk et al., 1990; Jason et al., 1984; Scott et al., 2016; Xia et al., 2020), and also individual variability in head size (Colby et al., 2022), which may have affected EEG electrode positioning and the recorded brain areas. Throughout the delay period, significant beta-band coherence was observed between the LPFC and EEG over the occipitoparietal cortex. Unlike the continuous memory-dependent interareal interactions reported between LPFC and parietal regions in previous studies (Salazar et al., 2012; Antzoulatos and Miller, 2016; Jacob et al., 2018), we found transient increase in memory-dependent coherence around the anticipated distractor time, specifically when the monkeys remembered contralateral items (Figure 12). This difference may be attributed to the spatial extent of the beta-band activity, as LFP reflects activity captured over a smaller brain volume compared to EEG (Buzsáki et al., 2012). Furthermore, some previous studies have linked beta-band modulation in anticipation of stimulus (Van Ede et al., 2011; Liang et al., 2002; Gross et al., 2006). This transient beta-band coherence may suggest temporal orienting of the anticipated distractor as a mechanism to mitigate its deteriorating effect.

Behaviorally, the distractor did not reduce the monkeys’ rate of task completion but did induce trial-wise errors, reflected in increased saccadic endpoint variability and response times, similar to previous findings in primates (Brown, 1958; Reinhart et al., 2012; Suzuki and Gottlieb, 2013; Oberauer et al., 2018; Hallenbeck et al., 2021; for review, see Lorenc et al., 2021). However, unlike some of these studies, we did not find systematic relationships between our behavioral and neural metrics (Supplementary Figures 9 and 10). A key distinction is that these prior studies utilized distractor stimuli that shared the same feature space as target stimuli (e.g., both target and distractor stimuli are circular and spatially informative), which can bias decision towards the distractor (also known as congruency effect). In contrast, our task employed a checkerboard stimulus that spanned much of the visual field and was spatially unrelated to the task, not sharing the same feature space, thereby the feature of the distractor minimally biases decision making at the end of the delay period of working memory task. This is important because it implies that the increased decoding of memory items in our decoding analysis was not a result of systematic behavioral biases towards a distractor.

In sum, we demonstrated that theta- and beta-band oscillations across spatial scales in monkeys reflect anticipatory processes to a potential distractor during working memory maintenance. Overall, our findings highlight that interareal beta-band oscillations may facilitate brain-wide memory reactivation in the theta band, giving indirect support to previous proposals such that the beta-band synchronization reactivates latent memory representations (Spitzer and Haegens, 2017). On the other hand, as beta-band oscillations are linked to both inhibition and top-down control during working memory tasks (Miller et al., 2018; Lundqvist et al., 2024), our results may also suggest that the beta-band-based inhibition and disinhibition of theta-band oscillations during distractor anticipation induce replay-like activity across the brain, leading to memory-dependent activity after the anticipated distractor onset. Future research should explore the interaction between theta- and beta-band oscillations across brain regions during distractor anticipation, specifically to determine whether the observed memory replay reflects a conscious recall of information or a spontaneous recall-like process induced by inhibition and disinhibition of oscillatory activity.

