## Supplementary Figures for "Distractor anticipation during working memory is associated with theta and beta oscillations across spatial scales"

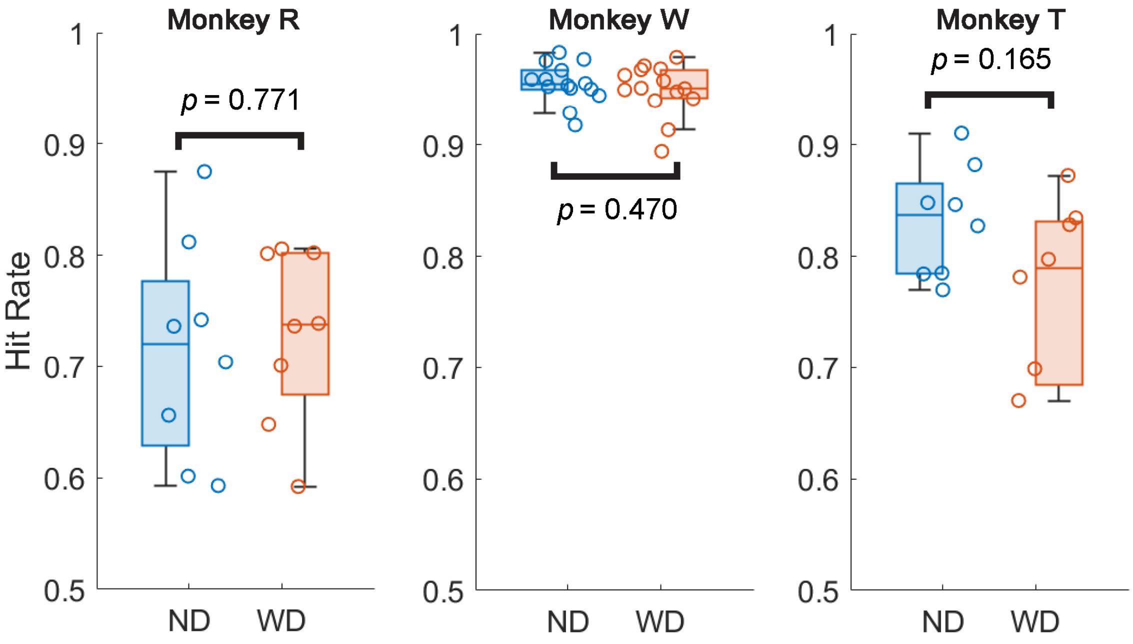


Supplementary Figure 1: Hit rate comparisons between ND and WD conditions showed no difference across monkeys.

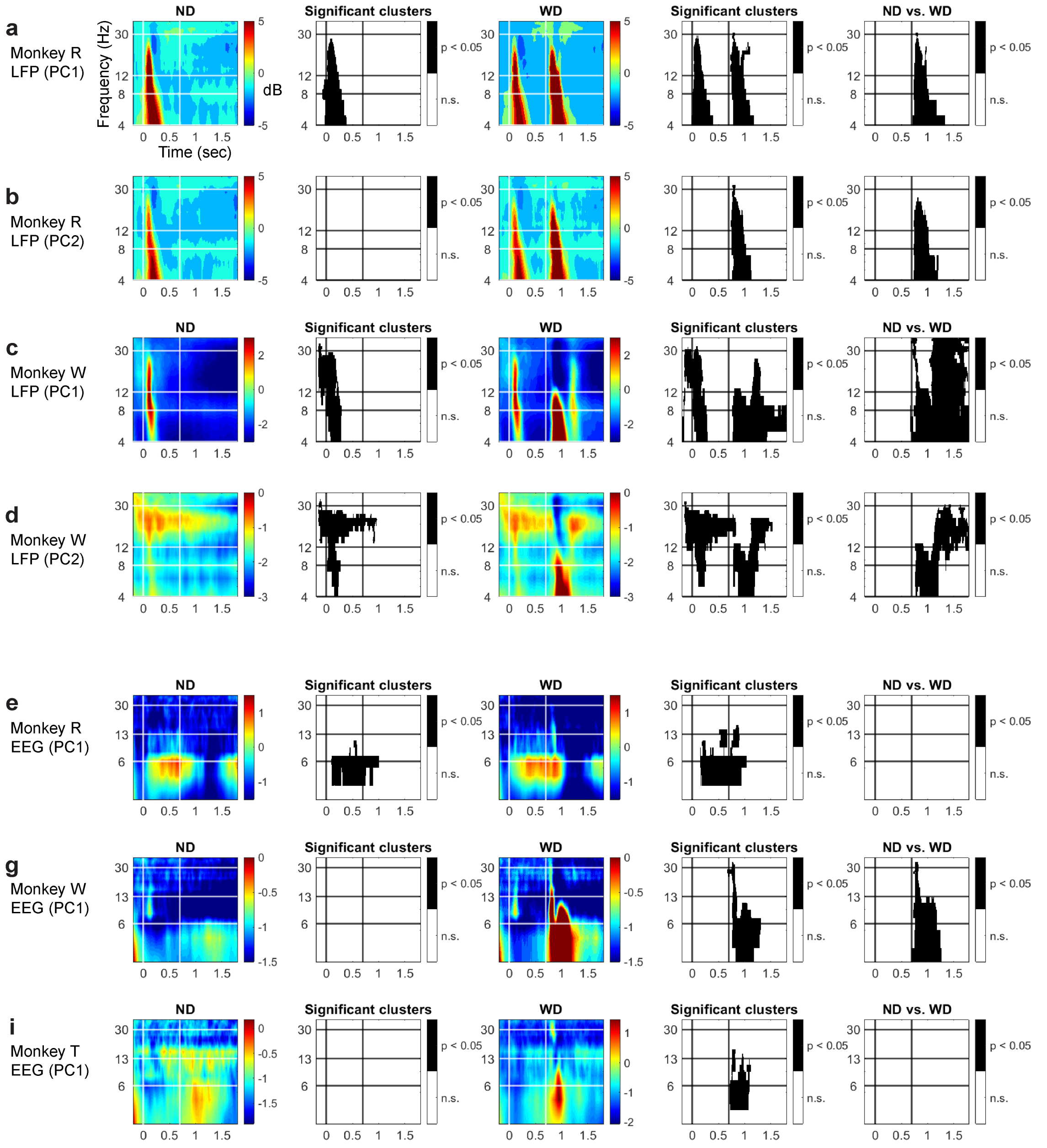


Supplementary Figure 2: Time-frequency representations of LFP PC1 (**a**, **b**) and PC2 (**b**, **d**) for monkeys R and W. Similarly, time-frequency representations for EEG PC1 for monkeys R, W, and T (**e**, **g**, and **i**, respectively). These representations, displayed in displayed in decibels (dB), are aligned to target onset (time = 0 ms), with the distractor onset indicated by a vertical line at 700 ms. Significant clusters ($\alpha$ = 0.05 and *cluster* $\alpha$ = 0.05) shown in black-and-white maps for each representation. The last column displays comparisons between the ND and WD conditions.

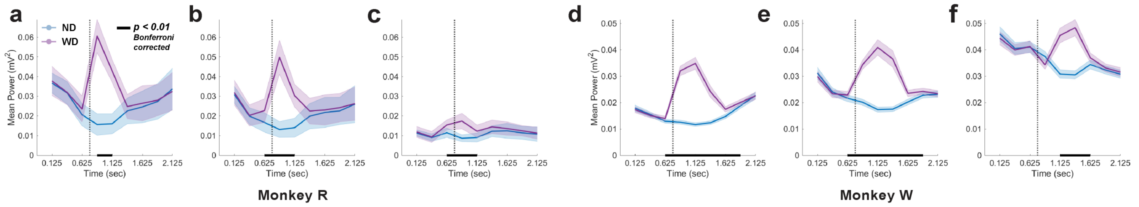


Supplementary Figure 3: Dimensionally reduced band powers (PC1) at theta (**a**, **d**), alpha (**b**, **e**), and beta (**c**, **f**) bands of the LPFC LFP collected from monkeys R and W, respectively. Horizontal black bars indicate significant time bins (500-ms bin) at which the difference between ND and WD conditions was observed ($p < 0.01$; Bonferroni correction applied across time bins; N = 9 bins)

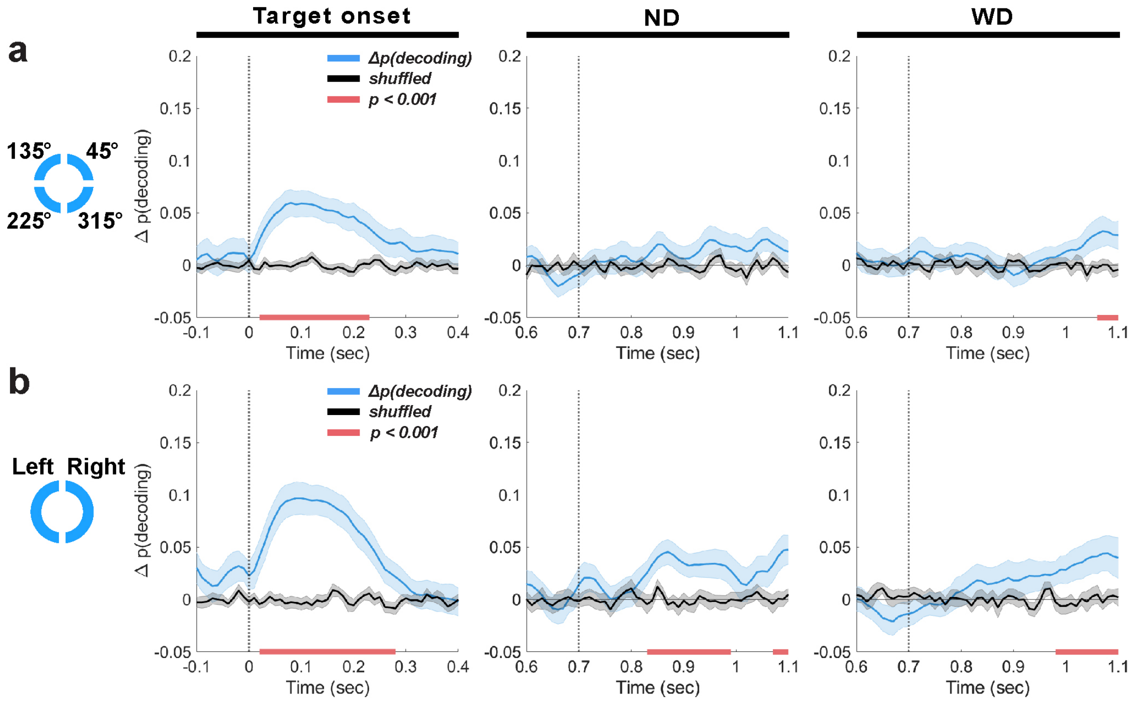


Supplementary Figure 4: Using EEG theta-band activity (1 - 6 Hz), decoding of location (**a**) or left-versus-right visual hemifield location (**b**) of a memory item from incorrectly performed trials by evaluating them on the classifiers trained on correct trials across all monkeys. Decoding results are aligned to target onset (time = 0 ms), anticipated distractor onset (ND; time = 700 ms), and actual distractor onset (WD; time = 700 ms). Mean decoding performance (blue traces, ±1 SEM) and trial-shuffled performance (black traces, ±1 SEM) are shown, with significant clusters ($p < 0.001$ for all trial conditions) indicated by horizontal red bars at the bottom. The results were obtained by evaluating classifier performance on incorrect trials after being trained on correct trials.

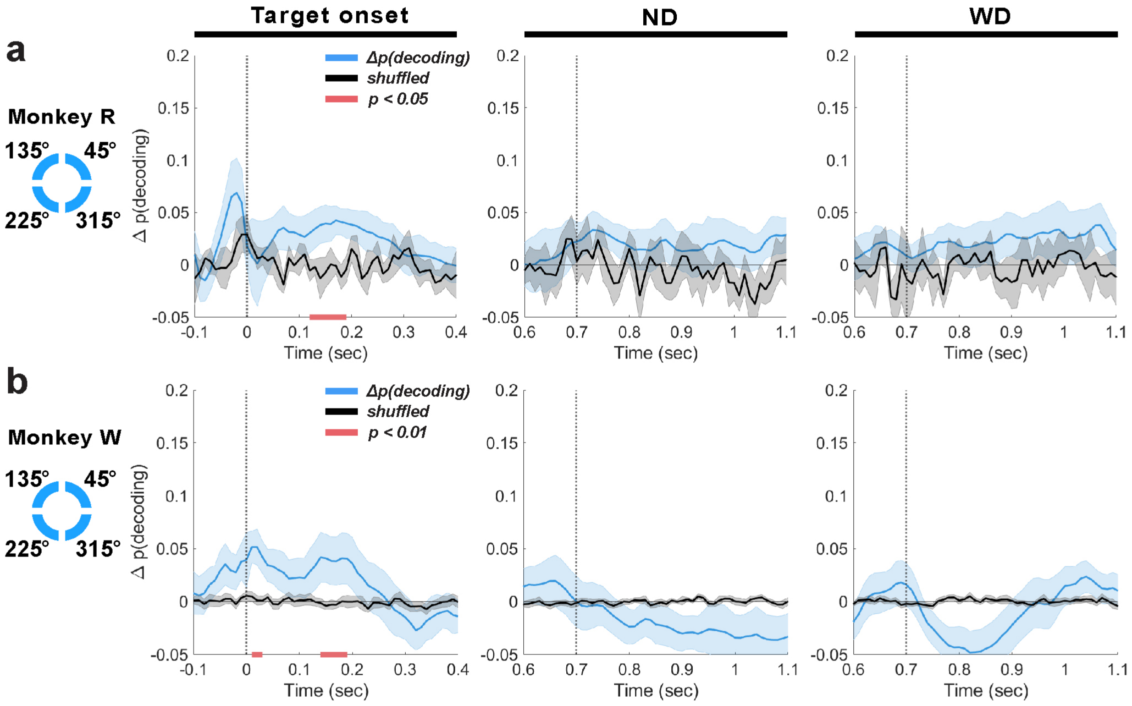


Supplementary Figure 5: Using LFP theta-band activity (4 - 8 Hz) from the LPFC, the location of memory items was decoded from incorrectly performed trials by testing these trials on classifiers trained with correct trials for monkeys R (**a**) and W (**b**). Decoding results are aligned to target visual stimulus onset (time = 0 ms), anticipated distractor onset (ND; time = 700 ms), and actual distractor onset (WD; time = 700 ms). Overall, weak and spurious decoding at target onset, implicating that the items were not encoded accurately from the start of the trial. No decoding available in both post-anticipated-distractor and post-distractor periods. Mean decoding performance (blue traces, $\pm 1$ SEM) and trial-shuffled performance (black traces, $\pm 1$ SEM) are shown, with significant clusters ($p < 0.05$ for monkey R and $p < 0.01$ for monkey W) indicated by horizontal red bars.

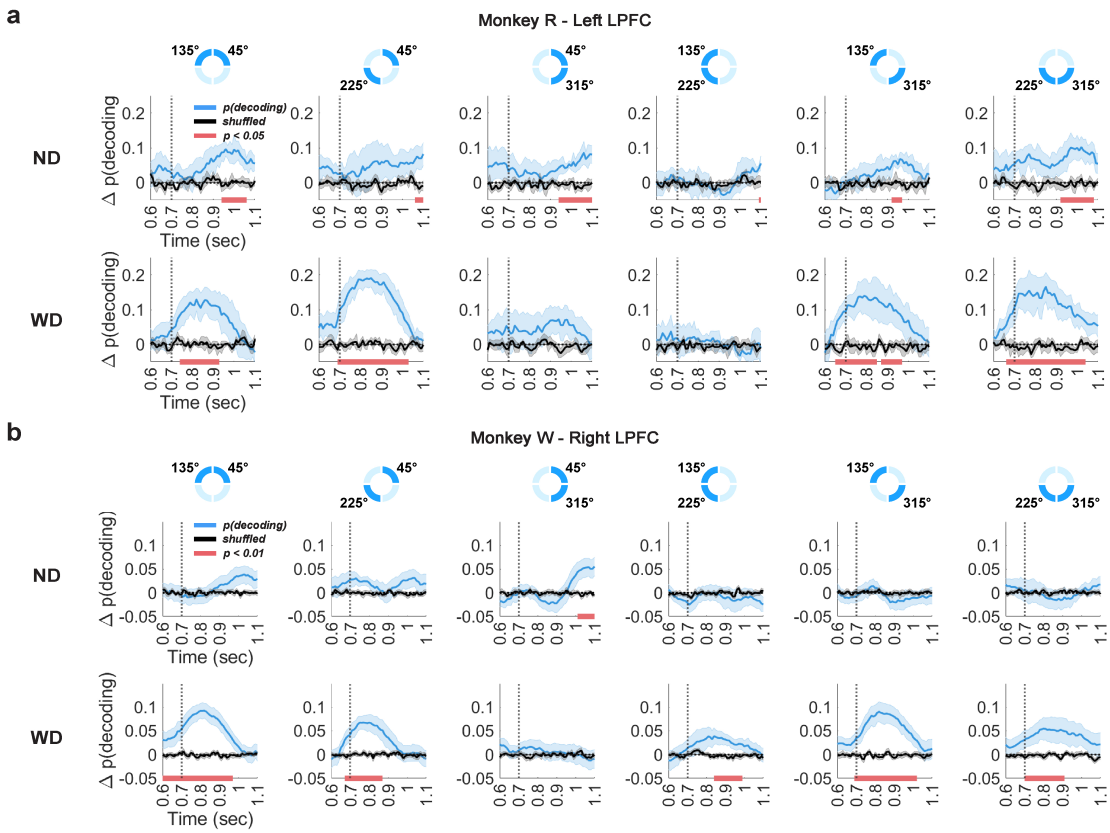


Supplementary Figure 6: Decoding results based on LFP theta-band activity (4 - 8 Hz) for all pairwise combinations of item locations (${}_4C_2 = 6$; 45° vs. 135°, 45° vs. 225°, 45° vs. 315°, 135° vs. 225°, 135° vs. 315°, and 225° vs. 315°) for monkey R (**a**) and monkey W (**b**), aligned to either the anticipated distractor onset (ND) or the actual distractor onset (WD) indicated by the vertical line at 700 ms. Mean decoding performance (blue traces, ±1 SEM) and its trial-shuffled version (black traces, ±1 SEM) are shown, with significant clusters ($p < 0.05$ for monkey R and $p < 0.01$ for monkey W) indicated by horizontal red bars.

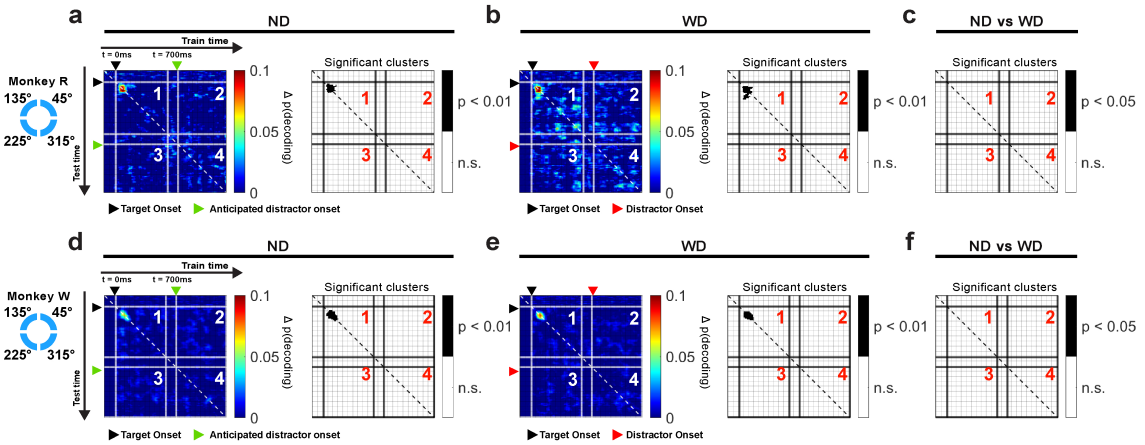


Supplementary Figure 7: Cross-temporal generalization using LFP beta-band activity (13 - 30 Hz) for monkeys R and W based on the locations of items for ND (**a** and **d**) and WD (**b** and **e**) conditions, with their differences (**c** and **f**). Panel 1 generalizes within the post-target time period (0 ms - 500 ms); panels 2 and 3 generalize between post-target onset (0 ms - 500 ms) and (anticipated) post-distractor onset time periods (700 ms - 1200 ms); panel 4 generalizes within the (anticipated) post-distractor onset time (700 ms - 1200 ms); significant clusters are shown in black-and-white maps. Black triangle: target onset; red triangle: distractor onset (WD); green triangle: anticipated distractor onset (ND).

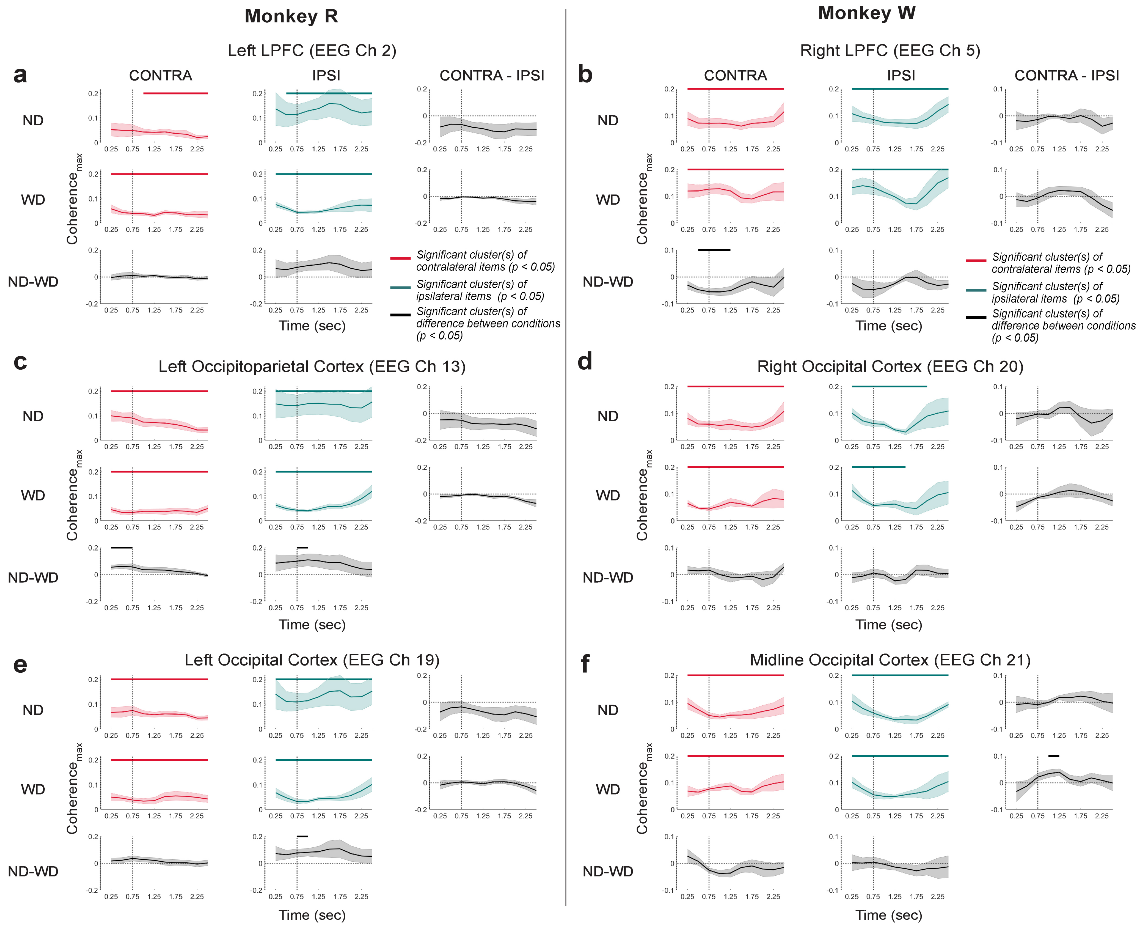


Supplementary Figure 8: Time-resolved beta-band coherence from incorrectly performed trials. Coherence estimates between the LPFC LFP and representative EEG channels, consistent with Figure 12. (**a**,**c**,**e**) coherence$_{\text{max}}$ ($\pm$1 SEM) for monkey R across trial and item conditions. (**b**,**d**,**f**) Same but for monkey W. CONTRA (in red traces) refers to trials where monkeys remembered items displayed in the visual hemifield contralateral to the LPFC implant site, while IPSI (in green traces) refers to trials with items displayed in the visual hemifield ipsilateral to the implant site. The vertical line at 700 ms marks the anticipated or actual distractor onset. Differences across conditions in black traces ($\pm$1 SEM). Significance indicated by horizontal bars (cluster-based permutation test result with $\alpha = 0.05$ and $cluster\ \alpha = 0.05$) with their respective colors.

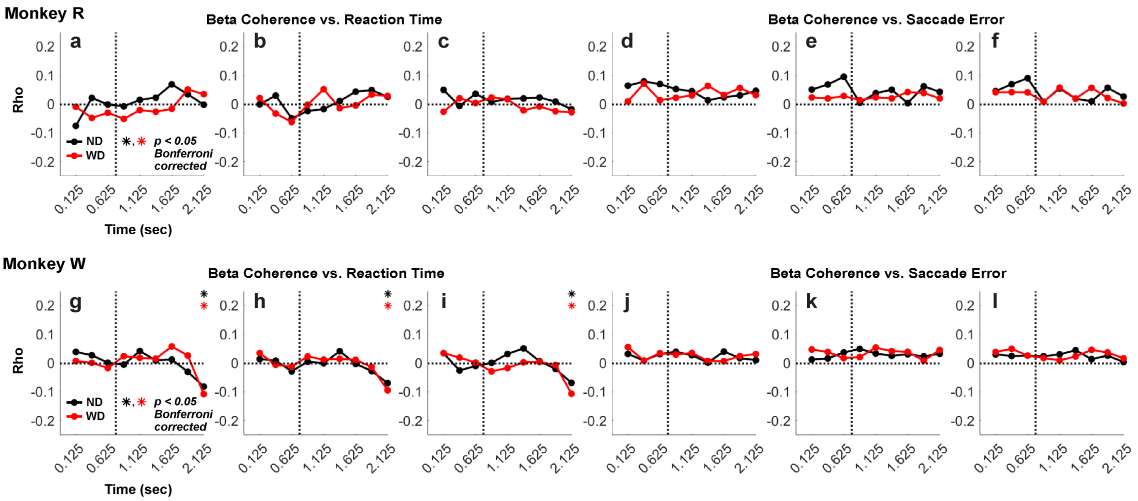


Supplementary Figure 9: Time-resolved correlation between beta-band coherence and behavioral error estimates –saccadic reaction time (in seconds) or saccade error (in radians). Here, trial-wise saccade error was calculated as the angular distance between saccade endpoints and their corresponding centroids. As described in methods, time-resolved coherence was estimated for each 500-ms window with a 250-ms overlapping window along time. For correlation, we used Spearman's rank correlation (Spearman, 1904) for saccadic reaction time and circular-linear correlation for saccade error (Fisher, 1995; Zar, 1999). Monkey R: Reaction time and saccade error correlated with the LFP-EEG beta coherence at channels 2 (**a**,**d**), 13 (**b**,**e**), and 15 (**c**,**f**). Monkey W: Reaction time and saccade error correlated with the LFP-EEG beta coherence at channels 5 (**g**,**j**), 16 (**h**,**k**), and 20 (**i**,**l**). The vertical line at 700 ms marks the anticipated or actual distractor onset. During the delay period (time $\leq$ 2000 ms), no correlation was achieved in all cases across monkeys. Asterisk ($*$): $\alpha < 0.05$, with ND in black traces and WD in red traces. Significance was Bonferroni-corrected for multiple comparisons across time bins (N = 9 bins).

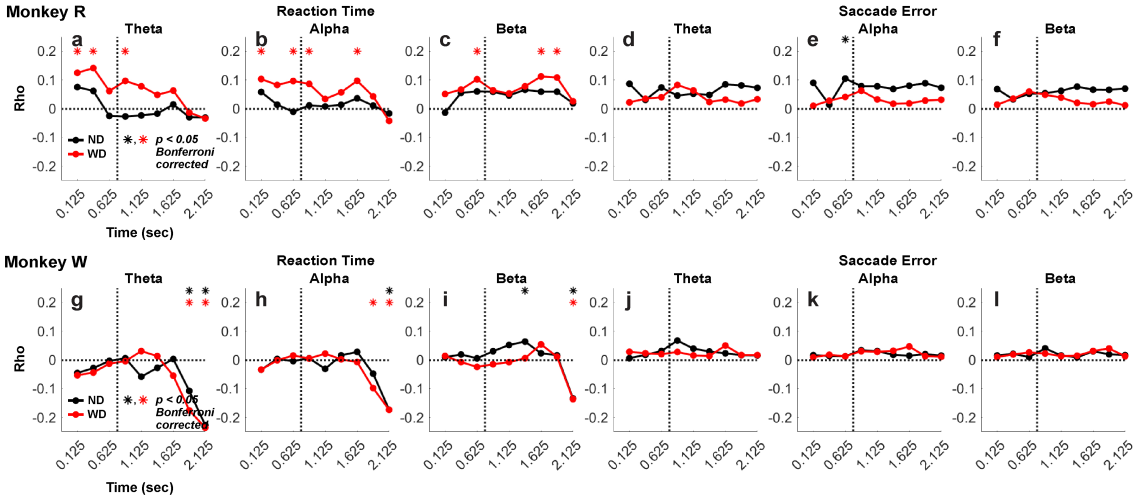


Supplementary Figure 10: Time-resolved correlation between band power and behavioral error estimates –saccadic reaction time or saccade error. Estimated mean band powers for theta (4 - 8 Hz), alpha (8 - 13 Hz) and beta (13 - 30 Hz) range from LFP PC1 data for each 500-ms window with a 250-ms overlapping window along time. Spearman's rank correlation for saccadic reaction time and circular-linear correlation for saccade error. Monkey R: Reaction time and saccade error correlated with LFP-EEG coherence at theta (**a**,**d**), alpha (**b**,**e**), and beta (**c**,**f**). Some correlation with the reaction time during the stimuli presentation was observed across bands. Monkey W: Reaction time and saccade error correlated with LFP-EEG coherence at theta (**g**,**j**), alpha (**h**,**k**), and beta (**i**,**l**). In monkey R, the correlation between power and reaction time appeared stronger in the trials with the distractor shown (WD). In monkey W, power only correlated with reaction time toward the end of the delay period (time $\geq$ 2000 ms). Overall, no correlation with saccade error was observed. The vertical line at 700 ms marks the anticipated or actual distractor onset. Asterisk ($*$): $\alpha < 0.05$, with ND in black traces and WD in red traces. Significance was Bonferroni-corrected for multiple comparison across time bins (N = 9 bins).
